# Dermonecrosis caused by spitting cobra snakebite results from toxin potentiation and is prevented by the repurposed drug varespladib

**DOI:** 10.1101/2023.07.20.549878

**Authors:** Keirah E. Bartlett, Steven R. Hall, Sean A. Rasmussen, Edouard Crittenden, Charlotte A. Dawson, Laura-Oana Albulescu, William Laprade, Robert A. Harrison, Anthony J. Saviola, Cassandra M. Modahl, Timothy P. Jenkins, Mark C. Wilkinson, José María Gutiérrez, Nicholas R. Casewell

## Abstract

Snakebite envenoming is a neglected tropical disease that causes substantial mortality and morbidity globally. The venom of African spitting cobras often causes permanent injury via tissue-destructive dermonecrosis at the bite site, which is ineffectively treated by current antivenoms. To address this therapeutic gap, we identified the aetiological venom toxins responsible for causing local dermonecrosis. While cytotoxic three-finger toxins were primarily responsible for causing spitting cobra cytotoxicity in cultured keratinocytes, their potentiation by phospholipases A_2_ toxins was essential to cause dermonecrosis *in vivo*. This evidence of probable toxin synergism suggests that a single toxin-family inhibiting drug could prevent local envenoming. We show that local injection with the repurposed phospholipase A_2_-inhibiting drug varespladib significantly prevents local tissue damage caused by several spitting cobra venoms in murine models of envenoming. Our findings therefore provide a new therapeutic strategy to more effectively prevent life-changing morbidity caused by snakebite in rural Africa.

**Significance Statement:** Spitting cobra venoms cause extensive local tissue damage surrounding the site of a snakebite. This damage cannot be effectively prevented with current antivenom treatments, and patients are often left with life-changing wounds. In this study we used cellular and mouse experiments to determine which toxins in African spitting cobra venom are responsible for causing tissue damage, revealing that a combination of two different types of toxins are required to cause pathology *in vivo*. We then showed that the repurposed drug, varespladib, which targets one of these toxin types, effectively prevents skin and muscle damage in mouse models of envenoming. Collectively these findings suggest that varespladib could be an effective new type of therapy for preventing snakebite morbidity in Africa.

## Introduction

Snakebite is a neglected tropical disease (NTD) that primarily affects rural communities in sub-Saharan Africa, South/South-east Asia and Latin America, and causes an estimated 138,000 deaths per annum, with a further 400,000 people maimed annually (*1*). Although historically receiving little attention, in 2017 the World Health Organization (WHO) added snakebite to their list of priority NTDs and subsequently devised a roadmap aiming to halve the number of deaths and disabilities attributed to snakebite by 2030 (*2*).

Snakebite patients affected by local tissue damage often require surgical tissue debridement or amputation to prevent the onset of life-threatening gangrene. These permanent sequelae greatly reduce the quality of life of most patients (*3*). Severe local pathology around the bite site results from cytotoxic, myotoxic and/or haemorrhagic venom toxins, and is most often observed after viper envenoming (*1, 4*). Whilst envenoming by most elapid snakes causes neurotoxic muscle paralysis and no local tissue damage, envenoming by several cobras (*Naja* spp.), most notably the African spitting cobras, causes little neurotoxicity but severe, rapidly developing swelling and tissue destruction that often leads to necrosis. These spitting cobra venoms also cause ophthalmia following defensive venom-spitting events (*5–7*). Spitting cobra bites are perhaps most frequent in sub-Sahel regions of Africa and include bites by *N. pallida* in eastern Africa (*8*), *N. mossambica* in southern Africa (*9*) and *N. nigricollis*, which has a wide distribution throughout northern parts of sub-Saharan Africa (*10*). Collectively, envenomings by spitting cobras substantially contribute to the numerous cases of severe local envenoming that result in permanent, life-afflicting morbidity across the African continent (*11*).

The cobra venom toxins predominantly associated with causing dermonecrotic pathology are the cytotoxic three-finger toxins (3FTx), hereafter referred to as CTx, which make up 56-85% of the total toxin abundance in spitting cobra venoms (*12*). CTx are well known to disrupt cell membranes and/or induce pore formation (*13–15*), which leads to cell death through a series of intracellular events related to the loss of control of plasma membrane permeability and via direct interaction with organelles, such as lysosomes (*13, 14*). Although CTx are the most abundant toxin type found in many cobra venoms (*12*), it is usually only those of the spitting cobras that cause severe local tissue damage after envenoming (*1*), suggesting that additional toxins are likely contributing to the severity of local envenoming. The next most abundant toxin family in several cobra venoms are the phospholipases A_2_ (PLA_2_). While the PLA_2_ toxins found in elapid venoms are often neurotoxic (*1*), cytolytic PLA_2_ also exist which can cause tissue necrosis (*16, 17*). For example, the spitting cobra PLA_2_ nigexine is cytolytic towards multiple tumour cell lines, and reduces cell viability and cell proliferation of epithelial human amnion cells (*18*). It has also been proposed that toxin combinations enhance venom cytotoxicity (*19–21*), with PLA_2_ toxins seemingly potentiating the effects of CTx (*21*). Understanding the relative contributions of different venom toxins to the severity of local envenoming is essential for the future design of targeted therapeutics to reduce the burden of snakebite morbidity – a key objective of our research.

Current treatment for snakebite envenoming relies on intravenous antivenom therapy, which consists of polyclonal antibodies generated via venom-immunisation of equines or ovines (*1*). While these therapeutics save countless lives, they are associated with several limitations that restrict their clinical utility, including: low affordability to those in greatest need (*1, 22*), limited efficacy against a breadth of snake species due to venom toxin variation (*22*), and high incidences of severe adverse reactions in the case of some antivenoms (*23, 24*). The need to deliver antivenom intravenously by a medical professional in a clinical environment prolongs the time from bite to treatment by an average of five to nine hours due to poor hospital-accessibility in the remote, rural tropical regions where most snakebites occur (*22, 25, 26*). Furthermore, intravenous antivenom antibodies are too large (typically ∼110 or ∼150 kDa) to rapidly penetrate the envenomed peripheral tissue and neutralise the aetiological cytotoxins – rendering antivenom treatment largely ineffective in reversing the swelling, blistering and necrotic outcomes of local envenoming (*1, 22, 23, 27, 28*). Collectively, these limitations highlight why the development of effective therapeutics is one of the core goals of the WHO’s roadmap to reduce the impact of snakebite envenoming (*2*).

To address these therapeutic gaps, in this study we used a combined approach of *in vitro* cell cytotoxicity assays and *in vivo* murine models to quantify and identify the toxins responsible for venom-induced dermonecrosis caused by the most medically important African spitting cobras. Our findings demonstrate that CTx are largely responsible for cytotoxic effects observed in cellular assays using human epidermal keratinocytes, but that PLA_2_ toxins contribute extensively to *in vivo* envenoming pathology by working in conjunction with CTx to cause dermonecrosis. Using the PLA_2_-inhibiting repurposed drug varespladib (LY315920) (*29–31*), we then demonstrate significant reductions in venom-induced dermonecrotic pathology *in vivo*, suggesting that the local injection of PLA_2_-inhibitory molecules following envenoming is a viable therapeutic strategy to reduce lifelong morbidity caused by spitting cobra snakebites.

## Results

### Spitting cobra venoms cause heterogenous dermonecrotic lesions *in vivo*

To define the local envenoming pathology caused by medically important cobras, we intradermally challenged mice with venom from African spitting cobras. Mice were injected with two different doses of venom from east (Tanzania) and west (Nigeria) African forms of the black-necked spitting cobra (*Naja nigricollis*), which collectively exhibit a broad distribution across sub-Saharan Africa and are known to induce severe local pathology in human victims (*6, 10*). Following euthanasia 72 hours (h) after venom challenge, the resulting dermonecrotic lesions were excised and analysed macroscopically and microscopically.

Macroscopically, the lesions were generally heterogenous in appearance, presenting with a dark-coloured necrotic centre surrounded by a ‘white’ area of tissue damage (**Fig. 1A**). To better define the lesion heterogeneity microscopically, we then performed histopathological analysis on haematoxylin and eosin (H&E) stained sections of the resulting lesions (**Fig. 1B, C** and **D**). Sections from control mice receiving PBS only presented the typical histological pattern of normal skin, including epidermis, dermis (with skin appendages), hypodermis, panniculus carnosus and adventitia (**Fig. 1B**). When areas of venom-induced skin damage were examined, there were clear histological differences between the macroscopically white and dark regions, with more pronounced damage observed in the latter. In samples collected from the dark lesions there was extensive damage to all layers of the skin. The epidermis was lost and a hyaline proteinaceous material was observed, while the dermis and hypodermis were severely damaged with skin appendages absent. Moreover, there was widespread muscle necrosis in the panniculus carnosus (**Fig. 1C**). In contrast, the white lesions were characterised by hyperplasia of the epidermis and an inflammatory infiltrate in the dermis, together with thrombi in some blood vessels; though in general, the structure of the various layers of the skin was preserved and the skin appendages were present (**Fig. 1D**). Thus, these two different macroscopic patterns of skin lesions correspond to different histopathological scenarios. Lastly, the areas of the dark and total lesions were measured, revealing a general trend towards dose-dependent increases in lesion size, and that the area of the dark lesion never exceeded half of the total lesion area (**Fig. 1E**).

**Fig. 1.**
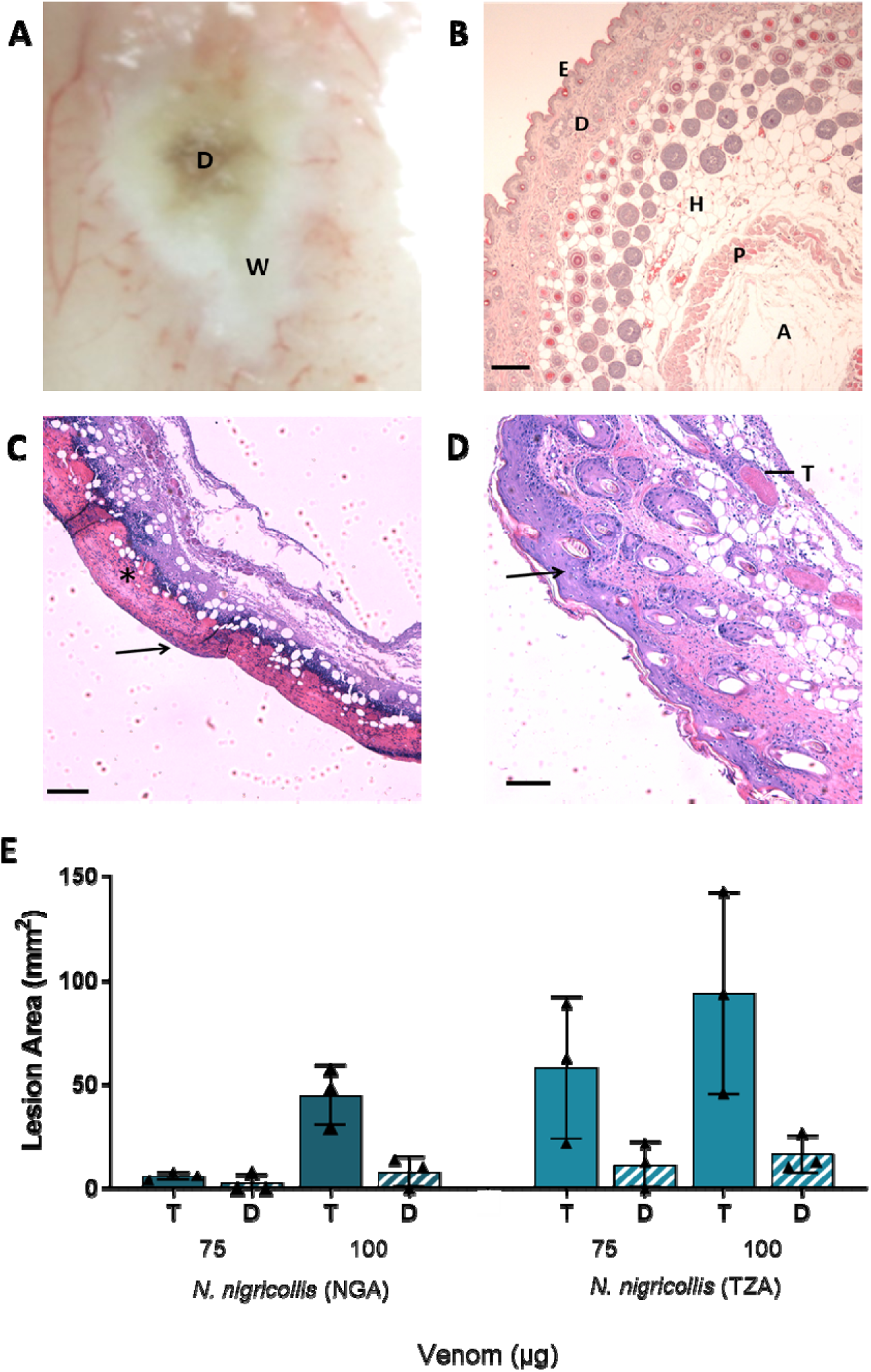
African spitting cobra venoms cause heterogenous dermonecrotic lesions *in vivo.* Groups of mice (n=3) were injected intradermally with two doses of spitting cobra venom and after 72 h the resulting lesions were excised for macroscopic quantification of damaged areas and histological assessment. **A**) Representative macroscopic image of a skin lesion induced by 100 µg of venom from West African (Nigeria) *N. nigricollis*, in which a dark central area (D) of necrosis is observed surrounded by a white area (W) of skin damage. **B-D**) Representative light micrographs of sections of the skin of mice injected with PBS or West African *N. nigricollis* venom. **B**) Skin injected with PBS showed a normal histological appearance including the epidermis (E), dermis (D), hypodermis (H), panniculus carnosus (P) and adventitia (A). **C**) Light micrograph of a section of skin corresponding to a dark area of venom-induced damage. All skin layers were affected, with loss of epidermis (arrow) and skin appendages in the dermis. A proteinaceous hyaline material was observed (*). **D**) Light micrograph of a section of the skin corresponding to a white area of damage from a mouse injected with venom. There was an increase in the thickness of epidermis (hyperplasia; arrow) and inflammatory infiltrate in the dermis. Thrombi (T) were observed in some blood vessels. **E**) The area of dermonecrotic lesions caused by *N. nigricollis* (West African, Nigeria [NGA]; East African, Tanzania [TZA]) venoms at different doses. Bars show the mean area of the total lesions (T) in comparison to the dark central areas (D) of greatest intensity, and error bars represent the standard deviation from the mean. Scale bars in B-D represent 100 µm.

### Venom CTx are predominately responsible for cytotoxic effects in cell culture

To identify which toxins in spitting cobra venoms are responsible for causing the dermonecrotic effects observed *in vivo*, we first identified and quantified the cytotoxic potency of venom constituents using cell cytotoxicity methods in immortalised human epidermal keratinocytes (HaCaT cell line). This work was performed using East African (Tanzania) *N. nigricollis* venom, which was separated into its distinct constituents via gel filtration and cation exchange chromatography, followed by further purification using hydrophobic interaction or hydroxyapatite chromatography (**Fig. S1-S9**). The identity of the isolated toxins was confirmed by mass spectrometric analysis (**Table S1**). The HaCaT cells were then exposed to either crude venom, four purified CTx (CTx1 [UniProt: P01468], CTx1v [UniProt: P01468], CTx3 [UniProt: P0DSN1] and CTx4 [UniProt: P01452]), or two purified PLA_2_ (basic [UniProt: P00605] and acidic [UniProt: P00602]), and combinations consisting of all CTx (at a ratio reflective of relative abundance in the venom; ∼3:1:1:1), all PLA_2_ (1:1), and all CTx and PLA_2_ together in a 2:1 ratio (reflective of that found in the crude venom (*32*)). Following venom exposure we performed thiazole blue tetrazolium (MTT) assays to assess cell viability via measures of metabolic activity (*33, 34*) multiplexed with propidium iodide (PI) assays as an indicator of cell death associated with plasma membrane disruption (*35*) (**Fig. 2**).

**Fig. 2.**
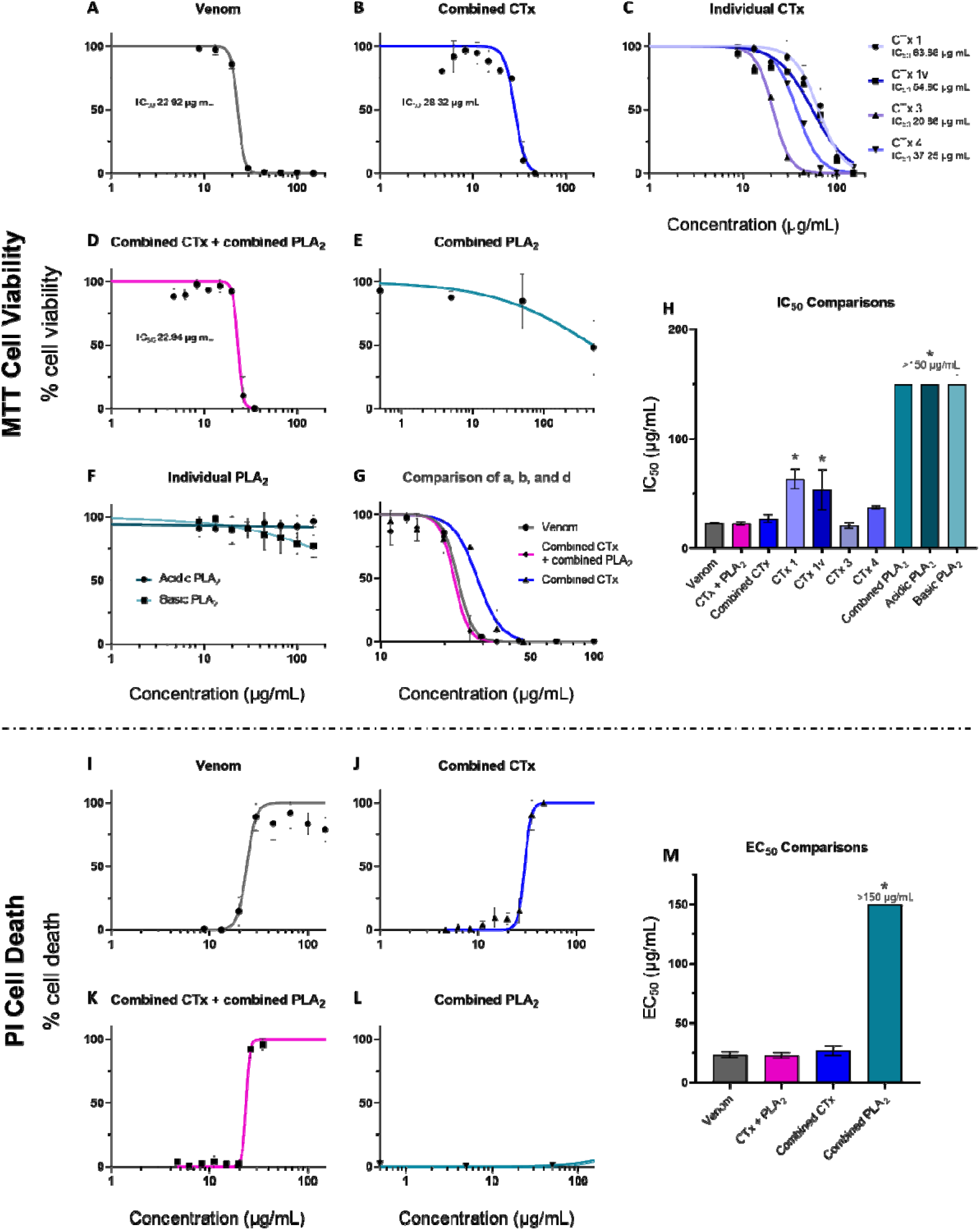
Crude venom and purified CTx inhibit cell viability, with CTx venom activity modestly potentiated by PLA_2_ toxins. Cell viability was measured in immortalised human keratinocytes (HaCaT cells) using MTT assays (A-H) and multiplexed with propidium iodide (PI) assays to measure cell death (I-M). HaCaT cells were treated for 24 h with serial dilutions of East African (Tanzania) *N. nigricollis* venom or its isolated toxins. MTT concentration-response curves are shown for: **A**) crude east African *N. nigricollis* venom, **B**) combined purified CTx, **C**) individual purified CTx, **D**) combined CTx and combined PLA_2_ together, **E**) combined purified PLA_2_, and **F**) individual purified PLA_2_. **G**) Direct comparison of the concentration-response curves caused by crude venom, combined CTx, and the combined CTx + combined PLA_2_ together. Note the different scale on the x-axis in comparison with panels A-F. **H**) IC_50_ value summary of the various venom toxins displayed in A-F using MTT assays. PI concentration-response curves are shown for: **I**) crude *N. nigricollis* venom, **J**) combined purified CTx, **K**) combined CTx and combined PLA_2_ together, and **L**) combined purified PLA_2_. **M**) EC_50_ values of the various venom toxins displayed in I-L using PI assays. For panels A-F, the data shown represent mean % cell viability and corresponding standard deviations. For panels I-L, the data shown represent mean % cell death and corresponding standard deviations. All data displayed are from three independent experiments with each condition conducted in triplicate. Data were normalised to 0-100% between the lowest and highest read values for analysis, then plotted as concentration-response curves using GraphPad Prism 9. For panels H and M, statistically significant differences determined by one-way ANOVAs with Dunnett’s multiple comparisons post-hoc tests, are denoted by asterisks: * (*P*<0.05).

MTT measurements of cell viability, taken after 24 h of treatment, showed that crude venom potently reduced cell viability (IC_50_ 22.9 μg/mL ± 0.7; **Fig. 2A**). Similarly, all four of the purified venom CTx reduced cell viability, with CTx3 being the most potent (IC_50_ 20.8 μg/mL ± 2.5), followed by CTx4 (IC_50_ 37.2 μg/mL ± 1.2), and then CTx1 and CTx1v, which showed similar potencies (IC_50_ 63.4 μg/mL ± 9.0 and 53.5 μg/mL ± 18.4, respectively) and were significantly less potent than CTx3 (*P* = 0.004 and *P* = 0.020, respectively; **Fig. 2C, H**). While the basic PLA_2_ visibly showed some cell viability inhibitory effects at the highest concentrations tested (≥100 μg/mL), neither of the two purified PLA_2_ alone (**Fig. 2F**) or combined in a 1:1 ratio (**Fig. 2E**) were sufficiently toxic to the cells to allow for the calculation of IC_50_ values. The combination of the four purified CTx gave a complete concentration-response curve with a resulting IC_50_ value approaching those obtained with crude venom (27.0 μg/mL ± 3.6 vs 22.9 μg/mL ± 0.7, respectively; **Fig. 2B**), though remained slightly right-shifted in comparison (**Fig. 2G)**. When the CTx and PLA_2_ combinations were pooled together in a 2:1 ratio, reflective of their toxin abundance in crude east African *N. nigricollis* venom (*32*), the resulting concentration-response curve became indiscernible from that of the crude venom and resulted in a near-identical IC_50_ value (22.6 μg/mL ± 1.5 vs 22.9 μg/mL ± 0.7, respectively) (**Fig. 2H**).

PI measurements of cell death taken 24 h post-treatment with the various toxin combinations and east African *N. nigricollis* venom showed similar patterns (**Fig. 2I-M**). The crude venom displayed potent cytotoxic effects resulting in EC_50_ values of 23.4 μg/mL ± 2.4, while the PLA_2_ combination did not cause sufficient cell death at the highest concentrations tested to calculate an EC_50_ value (**Fig. 2I, L, M**). The CTx combination resulted in an EC_50_ value close to that of the crude venom (EC_50_ of 26.8 μg/mL ± 4.1 vs 23.4 μg/mL ± 2.4, respectively), while the 2:1 ratio of CTx:PLA_2_ combinations together modestly decreased the EC_50_ value (22.8 μg/mL ± 2.4), but to levels highly comparable to those obtained with crude venom (**Fig. 2J, K, M**).

### The combination of purified CTx and PLA_2_ induces venom-induced dermonecrosis *in vivo*

To understand whether CTx are also predominately responsible for dermonecrotic venom activity *in vivo*, we performed comparative experiments in our murine preclinical model of envenoming (*36*). The minimum necrotic dose (MND) of East African *N. nigricollis* venom, i.e., the dose that induces a lesion in the skin of 5 mm diameter 72 h after injection (*36*), was determined to be 63 µg/mouse, and doses of purified CTx, PLA_2_, and CTx + PLA_2_ that reflect their relative mass contribution to the total crude venom protein were determined (CTx and PLA_2_ comprise approximately 60% and 26% by weight of *N. nigricollis* venom, respectively (*16, 32, 37–39*)). Thus, groups of mice received intradermal injections of either 63 μg of crude venom, 37.8 μg of the purified CTx combination, 16.4 μg of the purified PLA_2_ combination, or 37.8 μg plus 16.4 μg of the purified CTx and the PLA_2_ combinations, respectively, combined. After 72 h, animals were euthanised, dermonecrotic lesions excised, measured, photographed and processed for histopathological analysis (images of the resulting dermonecrotic lesions are presented in **Table. S2**).

Mean lesion areas resulting from venom injection were large and varied extensively among the experimental animals (52.0 mm^2^ ± 24.6) (**Fig. 3A**). Mice receiving the CTx and PLA_2_ combinations together (CTx + PLA_2_) developed lesions that did not differ significantly in size from those induced by the whole venom (27.7 mm^2^ ± 27.8; *P*>0.05), suggesting that these two groups of toxins are collectively responsible for recapitulating the major effects of the crude venom. In contrast with our cytotoxicity data, however, the CTx combination alone resulted in negligible dermonecrosis *in vivo*, with only one of the four experimental animals displaying a visible lesion, and the mean lesion size (2.1 mm^2^ ± 4.1) being significantly lower than that caused by the crude venom (*P* = 0.027). Again in contrast to the cell data, the PLA_2_ toxin combination resulted in three of the four mice developing visible lesions (8.0 mm^2^ ± 9.3), though the mean lesion size remained more than threefold lower than that observed with the crude venom and the CTx and PLA_2_ combination together. The overall severity of the lesions was also assessed using our recently developed, AI-based dermonecrosis quantification tool, VIDAL, which standardises and quantifies lesion size and intensity to calculate an overall dermonecrosis score in Dermonecrosis Units (DnU) (**Fig. S11**) (*40*). Quantification by VIDAL largely recapitulated the results above (**Fig. 3B**), confirming that lesion severity caused by crude venom (120.6 dermonecrotic units [DnU] ± 61.6) was not significantly different from that of the CTx + PLA_2_ combination (66.3 DnU ± 70.2), and that lesions caused by the CTx combination resulted in significantly less dermonecrosis (6.8 DnU ± 13.7, *P*=0.025) than the crude venom. Lastly, total dermonecrosis scores (*41*) were calculated after histopathological assessment of H&E-stained lesion cross-sections for each animal (**Fig. 3C**). These data revealed that mice injected with crude venom showed extensive damage in all layers of the skin (dermonecrosis severity score of 2.6 ± 1.6). The PLA_2_ treated mice exhibited the next highest dermonecrosis severity scores (2.0 ± 1.5), followed by those receiving the CTx + PLA_2_ combination (0.9 ± 0.8) and then the CTx combination only (0.3 ± 0.3). These latter two groups exhibited dermonecrosis severity scores that were significantly lower than that of the crude venom (*P*=0.0218 and *P*=0091, respectively). Scoring for individual skin layers can be seen in **Fig. S10**.

**Fig. 3.**
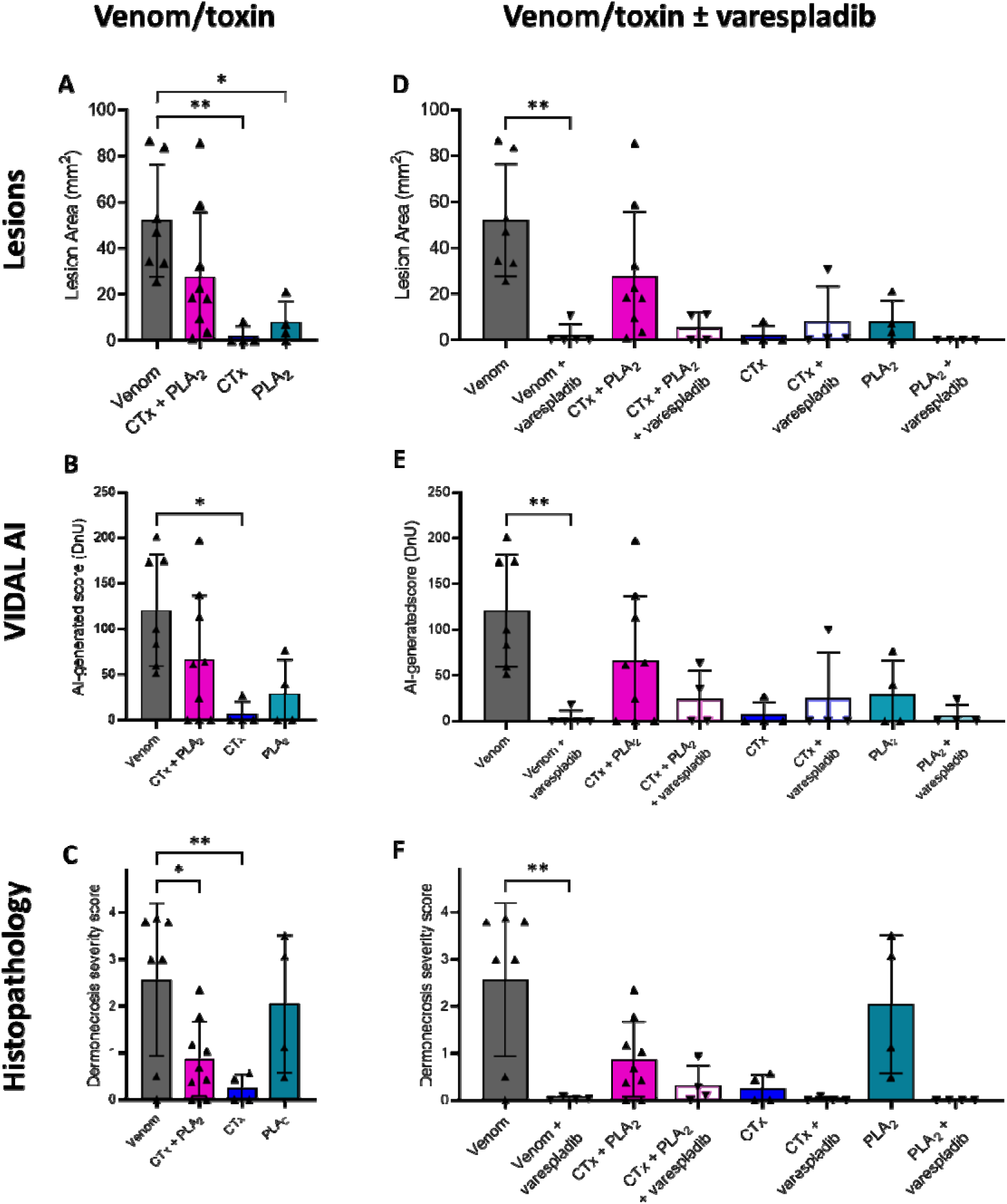
Spitting cobra venom causes dermonecrosis *in vivo* via CTx and PLA_2_ toxin potentiation, and inhibition of PLA_2_ toxins with varespladib reduces dermonecrosis severity. Groups of mice (n≥4) were intradermally injected with either East African (Tanzania) *N. nigricollis* venom or purified venom constituents (CTx, PLA_2_, or a combination of CTx + PLA_2_) at doses reflecting their relative abundance in crude venom, with or without the PLA_2_-inhibiting small molecule drug varespladib (19 μg). At 72 h post-injection, lesions were excised and examined macroscopically and histopathologically. **A**) A combination of venom CTx and PLA_2_ was required to recapitulate the dermonecrotic activity of crude *N. nigricollis* venom, as CTx and PLA_2_ toxins alone did not cause extensive dermonecrosis, as quantified via **A**) calliper-measurements of lesion height and width, and **B**) the lesion severity measuring AI tool, VIDAL. **C**) Histopathological analysis of excised lesions showed similar results, except for a more severe dermonecrotic effect of the PLA_2_ alone. Pre-incubation with the PLA_2_ inhibitor varespladib reduced dermonecrotic lesion severity caused by East African *N. nigricollis* venom with similar trends for CTx + PLA_2_ and PLA_2_, as quantified with **D**) callipers, **E**) VIDAL, and **F**) histopathological analysis. For damage scores of individual skin layers see **Fig. S10**. For panels A and D, the data shown represent mean lesion areas and corresponding standard deviations. Panels B and E show the mean lesion severity, as determined by VIDAL. Panels C and F show mean dermonecrosis severity scores and corresponding standard deviations calculated from those of the individual skin layers (**Fig. S10**). Statistically significant differences were determined by one-way ANOVAs followed by Tukey’s multiple comparisons post-hoc tests and are denoted by asterisks: * (*P*<0.05), ** (*P* <0.01). Error bars represent standard deviations.

### The PLA_2_ inhibitor varespladib protects against venom-induced dermonecrosis

Since our data demonstrated that venom-induced dermonecrosis relies on the combined effect of CTx and PLA_2_ venom toxins working together, we hypothesised that inhibiting just one of these toxin classes could significantly reduce the severity of venom-induced dermonecrosis *in vivo*. To that end, we repeated the experiments described above in the presence of the PLA_2_ inhibitor varespladib. Varespladib was originally designed for use in the treatment of cardiovascular diseases (*42–44*), but has recently entered phase II clinical trials for snakebite envenoming (*45*) following demonstration of its ability to prevent PLA_2_ toxin-driven systemic envenoming pathologies in animal models (*29, 30, 46, 47*).

We pre-incubated 19 μg of varespladib (*41*) with the same venom or purified toxin challenge doses before intradermally co-injecting mice and excising and analysing lesions 72 h later, as described above. The co-injection of varespladib with crude venom caused statistically significant reductions in lesion sizes from 52.0 mm^2^ (± 24.6) to 2.6 mm^2^ (± 5.3) (*P* = 0.001; **Fig. 3D**). Co-injection of varespladib with the CTx + PLA_2_ combination also caused a substantial reduction in mean lesion size from 27.7 mm^2^ (± 27.8) to 5.5 mm^2^ (± 6.4), although this reduction was not statistically significant. Unsurprisingly, varespladib did not affect the minor lesion formation observed in the group dosed with the CTx combination, though when varespladib was dosed alongside the purified PLA_2_, the resulting lesion sizes decreased from a mean of 8.0 mm^2^ (± 9.3) to no lesions being formed in any of the four experimental animals. These results were confirmed with the AI-generated dermonecrosis severity scores (**Fig. 3E**), from which the crude venom-induced lesions of 120.6 DnU ± 61.6 decreased significantly to 3.6 DnU ± 8.1 (*P=*0.0054) when co-incubated with varespladib. Similar effects of varespladib were seen when varespladib was co-incubated with the CTx + PLA_2_ combination (decreasing mean lesion score from 66.3 DnU ± 70.2 to 24.7 DnU ± 30.7) and PLA_2_ (decrease from 29.2 DnU ± 36.9 to 5.9 DnU ± 11.8), albeit these were not statistically significant. Decreases in lesion severity were not observed when varespladib was co-treated with CTx. Histopathological analyses of skin lesion cross-sections also correlated with the macroscopic assessment of dermonecrosis. Mice receiving venom pre-incubated with varespladib showed significantly less microscopic damage than those that received venom alone (dermonecrosis severity scores: venom, 2.6 ± 1.6; venom and varespladib, 0.0 ± 0.0; *P* = 0.0035) (**Fig. 3F**). Reductions in microscopic damage by varespladib were also observed in animals receiving either the PLA_2_ + CTx combination or PLA_2_ dose, although these were not statistically significant (dermonecrosis severity scores: PLA_2_ + CTx, 0.9 ± 0.8 vs PLA_2_ + CTx and varespladib, 0.3 ± 0.4; PLA_2_, 2.0 ± 1.5 vs PLA_2_ and varespladib, 0.0 ± 0.0). Full details of the dermonecrosis severity scores obtained across each individual skin layer are presented in **Fig. S10**.

To confirm that the inhibitory effect of varespladib is the sole result of inhibition of PLA_2-_driven toxicity, we performed MTT cell cytotoxicity assays, as described previously, using either East African *N. nigricollis* venom or the CTx combination preincubated with and without a cell-tolerated high dose of varespladib (128 μM) (*41*). As anticipated, while varespladib significantly reduced the cytotoxicity of crude venom (22.9 μg/mL ± 0.7 vs 30.3 μg/mL ± 2.0, *P* = 0.004), no significant effect on cell viability was observed when the PLA_2_ inhibitor was co-incubated with purified CTx (27.0 μg/mL ± 3.6 vs 29.2 μg/mL ± 3.8, *P*>0.05) (**Fig. 4**), thereby confirming that varespladib is only interacting with PLA_2_ toxins.

**Fig. 4.**
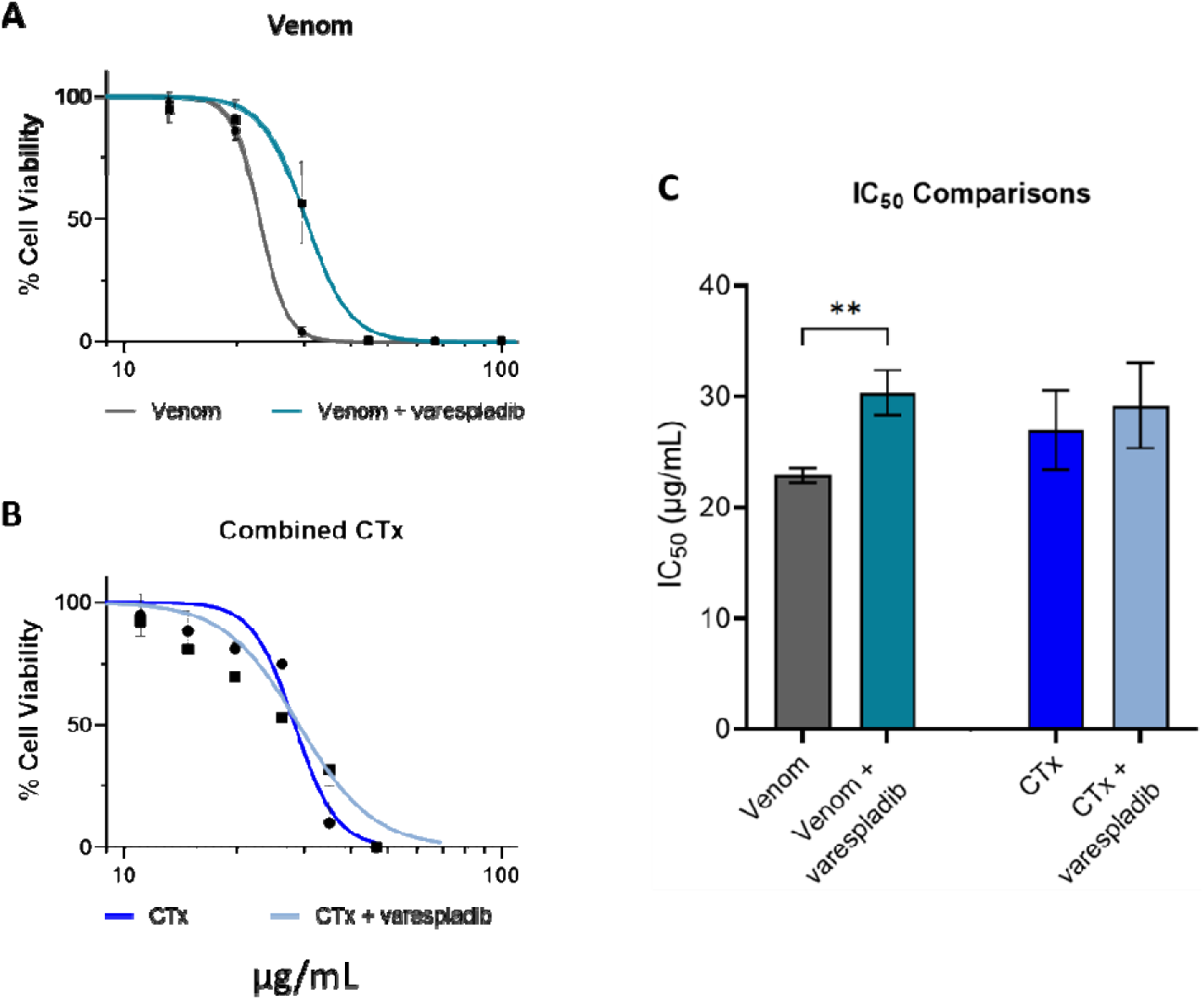
Preincubation with the PLA_2_ inhibitor varespladib has no effect on CTx-induced loss of cell viability in keratinocytes. Cell viability was measured in immortalised human keratinocytes (HaCaT cells) using MTT assays, with cells treated for 24 h with serial dilutions of (**A**) East African (Tanzania) *N. nigricollis* venom or (**B**) the purified CTx combination, with or without preincubation with 128 μM varespladib. The data shown represent mean percentage cell viability and corresponding standard deviations, with data normalised to 0-100% between the lowest and highest absorbance values for analysis, then plotted as dose-response curves using GraphPad Prism 9. All data displayed are from three independent experiments with each condition in triplicate. **C**) IC_50_ values of the venom and CTx combination, with and without varespladib, displayed in A and B. The data shown represent the mean IC_50_ values of curves and corresponding standard deviations. Statistically significant differences were determined by unpaired t-tests and are denoted by asterisks: ** (*P*<0.01).

To investigate whether the inhibitory effect of varespladib might extend to other cobra species, we next used venom from a related African spitting cobra species, the red spitting cobra *N. pallida*, which diverged from *N. nigricollis* around 6.7 million years ago (*21*), using the same *in vivo* model of dermonecrosis. Preincubation with varespladib resulted in complete abolition of lesion formation caused by *N. pallida* venom, with no lesions observed in any of the five experimental animals receiving the drug (venom, 32.4 mm^2^ ± 18.1 vs venom and varespladib, 0.0 mm^2^ ± 0.0; *P* = 0.0039) (**Fig. 5A**); a result that was also confirmed with the lesion severity scores calculated by VIDAL (59.0 DnU ± 28.2 versus 0.0 DnU ± 0.0, respectively; *P* = 0.0054) (**Fig. 5B**). Images of the resulting dermonecrotic lesions are presented in **Table. S3**. Further, histopathological assessment of venom-induced skin pathology also resulted in significant decreases in both total dermonecrosis scores (venom, 2.0 ± 1.2; venom and varespladib, 0.1 ± 0.1; *P* = 0.009) and for several individual skin layers analysed (epidermis, *P* = 0.003; hypodermis, *P* = 0.027; panniculus carnosus, *P* = 0.005) (**Fig. 5C** and **D**). Given that spitting cobra venom profiles share high levels of toxin similarity (*21, 32*), these findings provide confidence in the general effectiveness of varespladib against venom-induced dermonecrosis stimulated by geographically diverse African spitting cobra venoms.

**Fig. 5.**
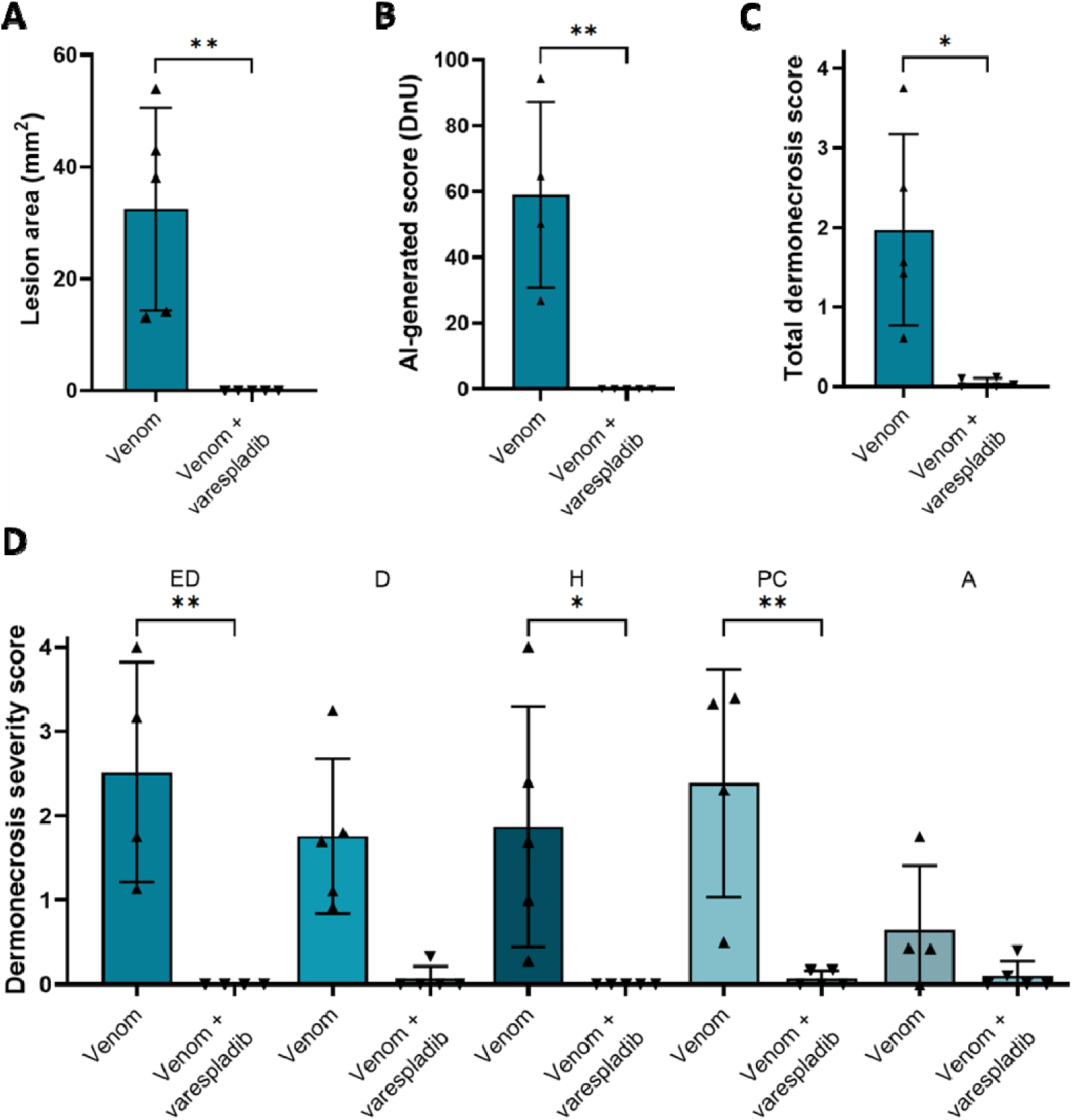
Dermonecrosis caused by *N. pallida* venom is prevented by the PLA_2_ inhibitor varespladib. Groups of mice (n=5) were intradermally injected with 25 µg *N. pallida* venom with or without the PLA_2_ inhibiting drug varespladib (19 μg). At 72 h post-injection, lesions were excised and examined, from which it was determined that the PLA_2_ inhibitor varespladib significantly reduced the size and severity of the dermonecrotic lesions caused by *N. pallida* venom as measured with **A**) callipers, **B**) VIDAL, and **C**) histopathological analysis of the **D**) different skin layers (ED, epidermis; D, dermis; H, hypodermis; PC, panniculus carnosus; A, adventitia). For **C** and **D**, the data shown represent the total mean dermonecrosis score of all layers, versus the mean damage score for each individual skin layer, respectively, and corresponding standard deviations. Statistically significant differences were determined by unpaired t-test comparisons for A, B, and C, and by two-way ANOVA, followed by Dunnett’s multiple comparisons tests for D. Statistically significant differences are denoted by asterisks: * = *P*<0.05, ** = *P*<0.01. Error bars represent standard deviations.

### Varespladib exhibits *in vivo* efficacy against spitting cobra induced dermonecrosis in delayed treatment models

Next, we sought to explore the inhibitory capability of varespladib in more biologically realistic models of snakebite envenoming, where treatment is delivered after venom challenge (*48, 49*). In addition, to further assess the cross-species and regional efficacy of varespladib we used another venom, this time from West African (Nigerian) *N. nigricollis*.

#### Dermonecrosis: Intradermal administration of varespladib

We repeated our previously described pre-incubation experiments and demonstrated again that the intradermal co-administration of venom with varespladib significantly reduced the size of the resulting skin lesions (42.8 mm^2^ ± 6.7 for venom vs 2.7 mm^2^ ± 6.5 for venom and varespladib; *P*<0.0001) (**Fig. 6A**). Then, to better understand whether varespladib could prevent dermonecrosis when administered after envenoming has occurred, and thus more accurately mimic a real-world snakebite scenario, we intradermally injected groups of mice with *N. nigricollis* venom (110 µg) followed by a second intradermal injection of varespladib (100 μg) in the same location at either 0, 15 or 60 minutes later. In all instances, we observed a significant reduction in the size of dermonecrotic lesions when varespladib was administered in comparison with the venom-only control (**Fig. 6B**). Reductions were most substantial in the group that received varespladib immediately after venom injection (0 min), where only one of the experimental animals presented with a small lesion 72 h later, resulting in significantly reduced mean lesion areas of 1.6 mm^2^ (± 2.7) compared with 34.0 mm^2^ (± 7.7) in the venom-only controls (*P*<0.0001) (**Fig. 6B**). While we observed reduced therapeutic potency with longer time delays between venom challenge and treatment, reductions in lesion sizes remained statistically significant at both 15- and 60-minutes post-venom challenge (13.6 mm^2^ ± 3.5 and 16.4 mm^2^ ± 5.5, respectively, vs 34.0 mm^2^ ± 7.7 with the venom only control; *P*<0.0001) (**Fig. 6B**).

**Fig. 6.**
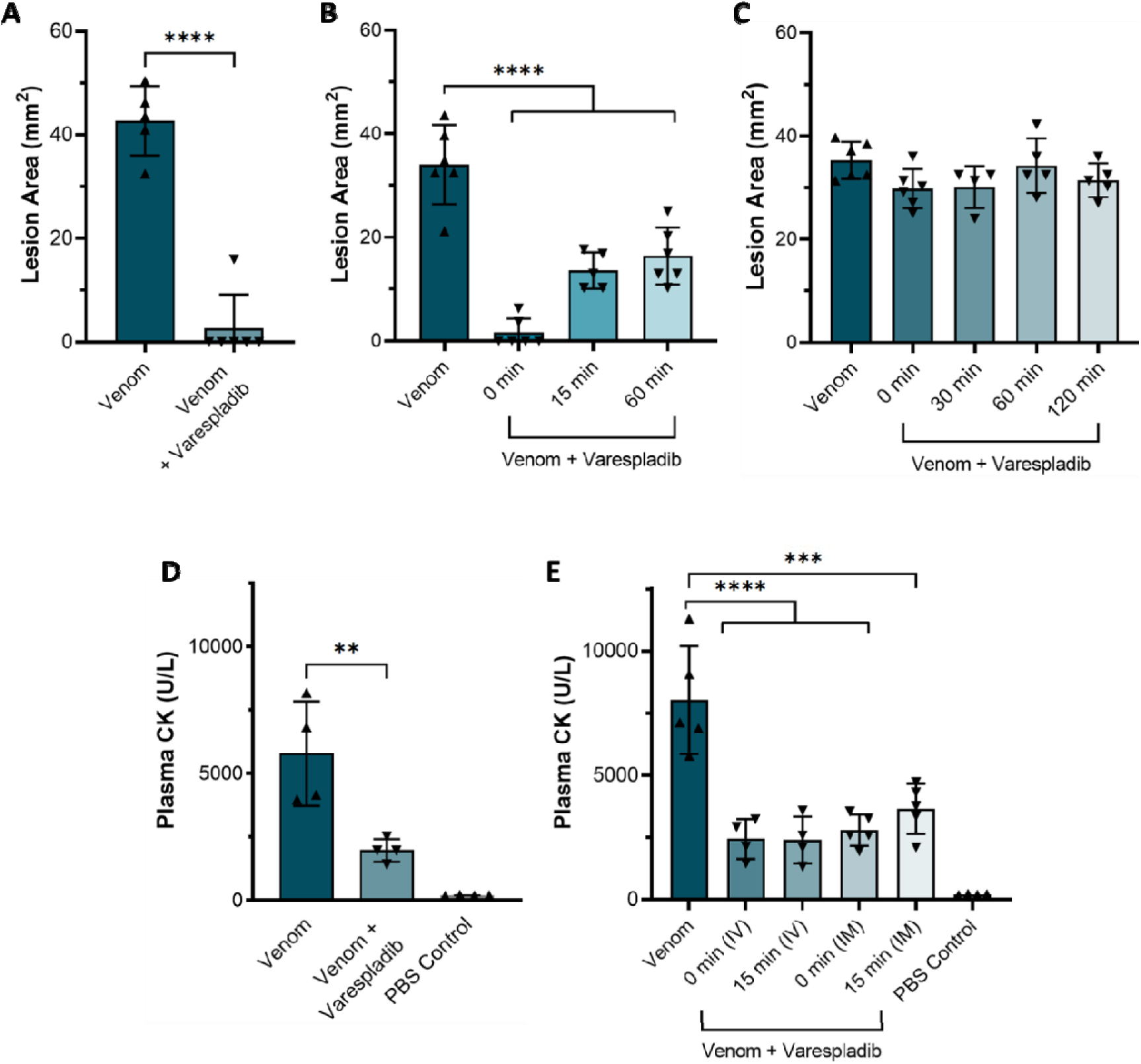
Delayed administration of varespladib following spitting cobra envenoming causes significant reductions in dermonecrosis and myotoxicity *in vivo*. Groups of mice (n≥4) were injected with either West African (Nigeria) *N. nigricollis* venom alone or followed by varespladib at a range of different timepoints and via different administration routes. **A**) Intradermal co-administration of preincubated venom (110 μg) and varespladib (20 µg) significantly reduced the size of skin lesions caused by west African *N. nigricollis* venom 72 h later. **B**) Intradermal administration of varespladib (100 µg) 0, 15 and 60 min after intradermal venom challenge (110 μg) resulted in significant reductions in the size of venom-induced dermonecrotic lesions. **C**) Intravenous delivery of varespladib (100 μg) at 0, 30, 60, and 120 minutes after intradermal venom challenge (110 μg) resulted in no protection against venom-induced dermonecrosis. **D**) Intramuscular delivery of varespladib (10 µg) co-incubated with 10 μg venom resulted in significant reductions in plasma CK activity induced by *N. nigricollis* venom. **E**) Intravenous (IV) and intramuscular (IM) delivery of varespladib (100 μg) 0 or 15 min after intramuscular venom challenge (10 μg) resulted in significant reductions in plasma CK activity induced by *N. nigricollis* venom. For panel A, statistically significant differences were determined by an unpaired t-test; for panels B-D, by one-way ANOVAs followed by Dunnett’s multiple comparisons post-hoc tests. Statistically significant differences are denoted by asterisks: * *P*<0.05, *** *P*<0.001, and **** P<0.0001.

#### Dermonecrosis: Intravenous administration of varespladib

To closer mimic current treatment of envenoming with antivenom, we next explored whether intravenous, rather than local, injection of varespladib could reduce venom-induced dermonecrosis. To that end, groups of experimental animals received the same intradermal dose of *N. nigricollis* venom (110 µg), followed by an intravenous injection of varespladib (100 μg) at either 0-, 30-, 60- or 120-min post-venom challenge. However, this dose of intravenous varespladib did not provide any reduction in the size of dermonecrotic lesions when compared with the venom-only control for any of the different dosing timepoints, including when varespladib was injected immediately after the venom challenge (35.3 mm^2^ ± 3.6 for the venom only control vs 30.1-34.3 mm^2^ for the various timepoint varespladib treatment groups) (**Fig. 6C**).

#### Myotoxicity: Intravenous and intramuscular administration of varespladib

Given the promising therapeutic findings observed when using varespladib against spitting cobra venoms that cause dermonecrosis, our final experiments assessed whether varespladib could also prevent venom-induced myotoxicity – an envenoming pathology also associated with the cytotoxic action of cobra venoms *in vivo* (*50–52*) and in cultures of myogenic cell lines (*16*). We induced myotoxicity in our murine envenoming model via intramuscular (gastrocnemius) venom injection and quantified muscle tissue damage by measuring plasma creatine kinase (CK) (*53*) activity 3 h later. When West African *N. nigricollis* venom (10 µg) was co-administered with varespladib (10 µg (*52*)), there was a significant reduction in resulting plasma CK levels compared to those obtained from mice receiving venom alone (venom, 5783.50 U/L ± 2055.86; venom and varespladib, 1969.50 U/L ± 448.67; *P* = 0.01) (**Fig. 6D**). Following this, we investigated whether the delayed administration of varespladib would retain efficacy against venom-induced myotoxicity, and we explored this via both intramuscular and intravenous delivery of the drug at the increased dose of 100 µg, matching the route of venom challenge and current antivenom delivery, respectively.

Treatment via both dosing routes resulted in significant reductions in venom-induced increases in plasma CK activity, irrespective of whether treatment was delivered immediately after venom challenge or 15 minutes later (54.48-69.59% reduction of plasma CK after varespladib injection vs venom-only control; *P*≤0.018 for all comparisons). Unlike that observed with the dermonecrosis experiments, there was little difference in drug efficacy between the two delivery routes tested, with the mean plasma CK activity of mice that received intravenous varespladib marginally lower than in those that received the therapy intramuscularly (2442.75 vs 2797.40, 0 mins; 2398.75 vs 3656.80, 15 mins; intravenous vs intramuscular, respectively), though these differences were not statistically significant (**Fig. 6E**).

## Discussion

Snakebite results in 2.5 million envenomings each year (*54*), yet outdated animal-derived antivenoms remain the only specific treatment available (*1*). Although these therapeutics are undoubtedly lifesaving interventions, their clinical utility is restricted by a lack of affordability, limited breadth of efficacy across different snake species, high incidences of adverse reactions associated with some products, and the necessity for snakebite patients to travel, often for several hours, to a hospital for intravenous antivenom administration (*1, 22*). Critically, antivenoms remain ineffective against the rapidly developing local pathology of snakebite envenoming, which in severe cases can lead to patients requiring tissue debridement around the bite site or amputation of the affected limb or digit (*22*). There is therefore an urgent and compelling need to develop new therapies against local envenoming by African and Asian cobras (*Naja* spp.), particularly African spitting cobras (*6, 55, 56*). Consequently, in this study we sought to: (i) characterise the dermonecrosis caused by the venoms of African spitting cobras, (ii) identify the primary aetiological dermonecrosis toxins found within these venoms and (iii) determine whether spitting cobra venom-induced dermonecrosis can be inhibited with the PLA_2_-inhibiting drug varespladib.

Using venoms from *N. nigricollis* (east and west African), we first characterised dermonecrotic pathology in a murine model of local envenoming. The results of these studies confirmed a dose-dependent relationship between the amount of venom injected and lesion severity, that the dermal lesions caused by spitting cobra venoms *in vivo* often contain a dark inner region surrounded by a lighter region of skin damage, and that these dark regions show more prominent microscopic damage than the lighter regions. Studies on dermonecrosis induced by cobra venoms, and their inhibition, should therefore consider this dichotomy in the assessment of the pathological effects of their venoms (*40*).

To elucidate the toxins responsible for inducing dermonecrosis we first used cell cytotoxicity assays with human epidermal keratinocytes as our model, from which we demonstrated that the CTx in east African *N. nigricollis* venom are the toxins predominately responsible for *in vitro* cytotoxicity. Despite PLA_2_ toxins having been described as having cytotoxic effects on different cell types (*18, 57*), the two types of PLA_2_ isolated from *N. nigricollis* venom had little effect on keratinocytes in isolation or when combined, though the basic PLA_2_ appeared to be slightly more cytotoxic at high concentrations than the acidic PLA_2_. Crucially, the combination of purified CTx did not completely replicate the cytotoxic potency of whole venom, which instead required using both the CTx and PLA_2_ combinations together (**Fig. 2G**), suggesting that PLA_2_ toxins at least mildly potentiate the cytotoxic activity of the CTx in cell culture experiments. This probable synergy between CTx and PLA_2_ has been documented in several previous studies that explored the venom of *Naja* and related elapid snake species (*19, 21, 58–61*); therefore, these findings are largely consistent with the literature, although the relative contribution of CTx versus PLA_2_-mediated cytotoxicity observed here is perhaps surprisingly skewed heavily towards CTx.

The likely synergy between the two toxin families became more apparent *in vivo* (**Fig. 3A-C**), where both CTx and PLA_2_ in isolation were found to only cause negligible dermonecrosis in mice, while their combination resulted in extensive dermonecrotic lesions approaching the size and severity of those formed by crude venom, notwithstanding wide inter-animal variability. Further, the CTx only caused modest damage to a single skin layer, the panniculus carnosus, in agreement with the known *in vivo* myotoxic effect of cytotoxic 3FTxs (*51*). Comparatively, venom and the CTx and PLA_2_ combinations together affected all skin layers, which is to be expected based on the large lesions formed by both *in vivo* (**Fig. S10**). A prior study by Rivel *et al.* also found that *N. nigricollis* venom caused murine necrosis of the dermis and loss of the epidermis, though they proposed that CTx were the primary driver for this damage (*62*). Similar conclusions were also made by Ho *et al.* when investigating dermonecrosis caused by the Asian non-spitting cobra *N. atra* (63). These findings contrast somewhat with the results of this study, which suggested that the purified CTx, which were predominately responsible for cytotoxicity in cell culture (**Fig. 2**), require the action of PLA_2_s to cause extensive dermonecrosis *in vivo*. Furthermore, histopathologically determined dermonecrosis severity scores in PLA_2_-induced lesions were high, despite the small gross lesion sizes. These data may suggest that PLA_2_ toxins can cause severe dermonecrosis focally, but without the presence of CTx this damage is limited in extent.

This necessity for both CTx and PLA_2_ toxins to be injected concurrently to cause extensive dermonecrosis is a notable finding, and directly correlates with recent data demonstrating that spitting cobra PLA_2_ toxins have evolved to potentiate the algesic effect of CTx to cause enhanced pain during defensive venom spitting (*21*). This means that spitting cobra dermonecrosis, and thus morbidity observed in snakebite victims, may be a direct consequence of the defensive origin of cobra venom spitting. In the context of snakebite therapeutics, our findings evidencing that a combination of CTx and PLA_2_ are required to cause dermonecrosis *in vivo* are notable, because they suggest that inhibiting either one of these toxin families could significantly reduce the overall pathology caused by the venom.

Varespladib inhibits PLA_2_ from a range of snake venoms, including non-spitting cobras, such as *N. naja*, *N. atra*, and *N. kaouthia* (*30*), other elapids (*31, 63*) and several viperids (*30, 31*). In contrast to PLA_2_, 3FTx are poorly immunogenic due to their small size and are non-enzymatic, making them more challenging therapeutic targets (*64*), and to date no broadly inhibitory anti-CTx repurposed drugs have been identified. Consequently, in this study we explored the therapeutic potential of varespladib, which has been selected as a lead candidate PLA_2_ inhibitor, and is currently undergoing clinical development for snakebite (*45*). However, research associated with varespladib has primarily focused on its potential utility in preventing or delaying the onset of systemic envenoming (*45, 46*). It has not, until now, been explored in the context of local necrosis following spitting cobra envenoming. Our *in vivo* preclinical efficacy experiments demonstrate that varespladib holds much therapeutic promise for this indication, as co-treatment with varespladib significantly inhibited the formation of dermal lesions caused by east and west African *N. nigricollis* and *N. pallida* venoms (**Fig. 3, 5,** and **6**). We found no evidence that varespladib inhibits the activity of CTx (**Fig. 3** and **4**), and thus these data support our hypothesis that a single drug targeting one toxin family (PLA_2_) can significantly reduce the severity of local envenoming caused by cobra venom. This is particularly noteworthy when considering that *Naja* venoms typically contain more CTx than PLA_2_, often twice as much based on venom weight, and that spitting cobra venoms share a relatively high degree of compositional similarity to one another (*12, 21, 32, 65*), particularly in the wider context of inter-specific venom variation (*12, 66*). Given that varespladib previously entered Phase III clinical trials for other indications (*67*), and that its oral prodrug form varespladib-methyl has entered Phase II clinical trials for snakebite in the USA and India (*45*), our findings suggest that repurposing this drug as a broad-spectrum treatment for preventing spitting cobra-induced dermonecrosis could be a valuable future application to mitigate snakebite morbidity in Africa.

Despite these exciting findings, it was important to address the limitations with the animal model of venom-induced dermonecrosis described above (*49*), where venom challenge and treatment are preincubated and co-administered in a manner artificial to real-world treatment of snakebite envenoming. Consequently, we assessed whether the observed efficacy of varespladib held when used as a treatment after venom challenge, for which we used west African *N. nigricollis* venom (**Fig. 6**). Independent intradermal injection of varespladib at the same site of the venom challenge resulted in significant reductions in the resulting venom-induced lesion sizes, even when treatment was delayed until 60 minutes after envenoming, and treatment with varespladib immediately after venom challenge resulted in comparable efficacy to when the drug was co-administered in the preincubation model (**Fig. 6A, B**). Together, these data suggest that varespladib introduced directly into the tissue where a victim was bitten could significantly reduce the resulting dermonecrosis, particularly if administered soon after a bite. Transdermal drug delivery systems are well-established approaches that could be readily applied here to achieve rapid delivery of varespladib to snakebite victims in a community setting (*68, 69*). Such an approach has the potential to drastically reduce the time between bite to initial treatment from hours or days (*25, 26, 70*) to minutes, thus drastically improving the prognosis of tropical snakebite victims.

Since varespladib administered intravenously proved ineffective against venom-induced dermonecrosis (**Fig. 6C**), these findings suggest that an intravenous, and therefore also an oral, version of the drug is less likely to be effective at preventing dermonecrosis. However, these data may simply reflect that, at the dose tested, insufficient varespladib is able to rapidly penetrate from the circulation into the affected peripheral tissue to prevent venom toxicity. Additionally, perhaps a different venom-inhibiting molecule with superior tissue-penetrating properties to varespladib would prove more effective in such experiments. Pharmacokinetic (PK) experiments are therefore required to robustly explore whether intravenous or oral delivery of PK-optimised doses of varespladib, or other inhibitors, might also be effective routes of delivery for the treatment of severe local dermonecrosis.

The myonecrosis-reducing effects of both intramuscular and intravenous injected varespladib (**Fig 6E**) suggests that both local and central methods of administration could be effective at preventing muscle toxicity associated with cobra snakebites. We hypothesise this difference in efficacy between myo- and dermo-necrosis rescue is due to the comparatively greater abundance of blood vessels in muscle versus cutaneous tissue, resulting in a greater and more rapid distribution of intravenously administered varespladib to the former (*71, 72*).

Despite the promising results of this study, there are several limitations. Firstly, the variation between the results of our *in vitro* and *in vivo* assays demonstrates that the action of spitting cobra venoms or toxins on keratinocytes does not replicate the complex pathological effects caused in skin *in vivo*. This highlights the need for further research into developing more accurate *in vitro* models of dermonecrosis, with organoids, organotypics and/or *ex vivo* skin models seeming likely to be valuable tools for future research (*73*). Additionally, murine models can only act as a guide for the effect a treatment may have in human patients (*74*), given differences between human and murine metabolism and immune systems (*75*), as well as differences in skin thickness and structure (*76*). Further, clinical trials will be needed to fully gauge the effect of varespladib against the local tissue-damaging effects of spitting cobra venoms in humans. Finally, our data was entirely focused on African spitting cobra venoms. Similar experiments should be performed using the venoms of cobra species from additional localities, particularly those found in Asia, to better determine the pan-cobra potential of varespladib as a novel treatment for local envenoming and to further investigate which toxins are primarily contributing to this pathology in other settings.

In summary, our study has shown that CTx found in spitting cobra venom are largely responsible for causing venom cytotoxicity in cellular assays, but that both CTx and PLA_2_ together are required to fully recapitulate the dermonecrotic effects of crude venom *in vivo*. Consequently, a drug that inhibits just one of these toxin types is likely to significantly reduce the overall dermonecrosis caused by crude venom. We tested this hypothesis using the repurposed PLA_2_-inhibiting drug varespladib and demonstrated impressive preclinical efficacy of the drug against three geographically diverse African spitting cobra venoms. Most notably, the local injection of varespladib was able to significantly reduce the extent of dermonecrosis, even when dosed up to an hour after venom challenge, and protection conferred by the drug also extended to venom-induced myotoxicity. Collectively, our data suggest that varespladib could become an invaluable new treatment against the tissue-damaging effects of spitting cobra venoms, which cause extensive morbidity in snakebite victims across the African continent.

## Materials and Methods

### Chemicals, Drugs and Biological Materials

See **Methods S1** for full detail on chemicals, drugs and biological materials used in this study.

### Venoms

Venom pools were from wild-caught animals of differing geographical origins, namely: *Naja nigricollis* (Tanzania and Nigeria; four individuals in each venom pool), and *N. pallida* (Tanzania; one individual). Venoms were sources and stored as described in **Methods S2**.

### Toxin isolation

Toxins from east African (Tanzanian) *N. nigricollis* venom were initially separated into 3FTx and PLA_2_ toxins using gel filtration chromatography (**Fig. S1-S3**). Acidic PLA_2_ consisted of toxins from two individual peaks (peaks 3 and 4) which were evaluated through SDS-PAGE and RP-HPLC, with peak 3 requiring a third chromatography step for full purity (**Figs. S4, S8; Table S1**). Basic PLA_2_ consisted of toxin from peak 11 and required dialysis before purity was confirmed through SDS-PAGE and RP-HPLC (**Figs. S5, S8; Table S1**). The 3FTx cytotoxins were found across a range of peaks (peaks 6, 8, 9, 12-14) and to be varying levels of purity as determined by SDS-PAGE and RP-HPLC, with some requiring further purification steps, including trypsin digestion and MS/MS analysis (**Figs. S7-S9**). See **Methods S3** for full details of the toxin isolation methods.

### Cells

The immortalised human epidermal keratinocyte line, HaCaT (*78, 79*), was purchased from Caltag Medsystems (Buckingham, UK). Cells were cultured in phenol red-containing DMEM with GlutaMAX supplemented with 10% FBS, 100 IU/mL penicillin, 250 µg/mL streptomycin, and 2 mM sodium pyruvate (standard medium; Gibco), per Caltag’s HaCaT protocol. For cell assays that contained the fluorescent dye PI, a medium formulated for fluorescence-based cell assays was used: FluoroBrite DMEM supplemented with 1% GlutaMAX 100x supplement, 1% FBS, 100 IU/mL penicillin, 250 µg/mL streptomycin, and 2 mM sodium pyruvate (minimally fluorescent medium; Gibco). The cells were split and growth medium changed 2x per week up to a maximum of 30 passages. Cells were maintained in a humidified, 95% air/5% CO_2_ atmosphere at 37 °C (standard conditions).

### Multiplexed MTT Cell Viability and PI Cell Death Assays

The MTT cell viability (*34*) and PI cell death assays were completed as described previously (*41*), with minor modifications. Briefly, HaCaT cells were seeded at 20,000 cells/well in black-sided/clear-bottomed 96-well plates in standard medium and incubated overnight in standard conditions. The following day, standard medium was aspirated and replaced with treatment solutions prepared in minimally fluorescent medium supplemented with 50 µg/mL PI: wells were treated in triplicate with 100 μL of crude venom (9.09-150.00 μg/mL), individual purified CTx (9.09-150.00 μg/mL), individual purified PLA_2_ (9.09-150.00 μg/mL), pooled purified CTx (5.96-44.67 μg/mL), pooled purified PLA_2_ (0.0005-500 μg/mL), or pooled purified CTx + PLA_2_ in a 2:1 ratio (4.68-35 μg/mL) sourced from east African (Tanzanian) *N. nigricollis* venom, then placed back in standard conditions for a further 24 hours. PI fluorescent readings (Ex_544_/Em_612_) were collected on a CLARIOstar Plus (BMG Labtech). The treatment solutions were then aspirated and replaced with MTT-containing minimally fluorescent medium (120 μL at 0.83 mg/mL), and the plates incubated for 45 min in standard conditions. Thereafter, the MTT-containing medium was aspirated, 100 μL DMSO added to each well to dissolve the formazan crystals, and absorbance (550 nm) was read for all wells on the CLARIOstar. Experiments were repeated independently on three occasions for each venom or toxin. Subsequently data were normalised to 0-100% between the lowest and highest absorbance values for analysis to represent %-cell death (PI) or %-cell viability (MTT), then plotted as dose-response curves using GraphPad Prism 9. IC_50_ (MTT) and EC_50_ (PI) values were calculated using the ‘[Inhibitor] vs. normalized response -- Variable slope’ and ‘[Agonist] vs. normalized response -- Variable slope’ functions, respectively.

### Animal ethics and maintenance

Animal experiment protocols were performed in accordance with ethical approvals from relevant bodies, using male SWISS (CD1) mice (18-27 g) housed in accordance with animal welfare standards. See **Methods S4** for full details on ethics and animal maintenance.

### *In vivo* dermonecrosis and co-treatment with varespladib using a preincubation model of envenoming

For initial experiments (**Fig. 1**) groups of mice (18-20 g; n=3) received intradermal injections in the ventral abdominal region of either 75 µg or 100 µg of the venoms of west African (Nigeria) or east African (Tanzania) *N. nigricollis*, dissolved in 50 µL of PBS; control mice were injected with 50 µL of PBS alone. After 72 h, mice were sacrificed by CO_2_ inhalation, the skins were removed, and the areas of the lesions on the inner side of the skin were measured. Then, skin samples were added to 3.7% formalin fixative solution and processed for embedding in paraffin. Sections (4 µm) were collected and stained with haematoxylin-eosin for microscopic assessment. To identify the toxins responsible for causing dermonecrosis and to assess whether varespladib inhibited this effect (**Figs. 3-5, 6A**), groups of mice (n=4-8) were briefly anaesthetised using inhalational isoflurane and then ID-injected in the rear flank with a 50 μL solution of either: (i) east African *N. nigricollis* venom (63 µg), (ii) corresponding proportional amounts of venom CTx (37.8 µg) or PLA_2_ (16.4 µg) isolated from the crude venom, (iii) these purified CTx and PLA_2_ combined in a 2:1 ratio, reflecting their relative abundance in crude venom (37.8 µg and 16.4 ug, respectively)(*16, 32, 38*), (iv) *N. pallida* (Tanzania) venom (25 µg), or (v) west African *N. nigricollis* venom (110 µg), all diluted in PBS (pH 7.4). The same experimental design was used for the varespladib-inhibition experiments except that every venom challenge dose was co-incubated with 19 μg of varespladib (*41*) (**Figs. 3-5**) or 20 μg of varespladib (**Fig. 6A**) (diluted in 98.48% pH 7.4 PBS and 1.52% DMSO) for 30 minutes at 37 °C and then kept on ice until shortly before intradermal injection. Following dosing, all experimental animals were observed frequently to ensure that no signs of systemic envenoming presented (e.g., neuromuscular paralysis), and the development of local lesions were monitored for 72 hrs. Thereafter, experimental animals were humanely euthanised via inhalational CO_2_, and the skins around the injection site dissected. The height and width of the lesions on the inner side of the skin were measured in two directions with digital calipers, placed on A4 printout sheets to standardise the AI-lesion analyser which are available in Jenkins, *et al.* (*79*), and photographed using a Sony DSC-W800 camera. Excised lesions were cut into cross-sections down the centre of the lesions, placed in plastic tissue cassettes (Sigma-Aldrich [Merck]; Z672122) and fixed in 3.7% formalin pots (CellPath; 13191184) for a maximum of 72 h prior to preparation for downstream histopathology.

### Delayed treatment models of *in vivo* dermonecrosis with varespladib

To assess inhibition of dermonecrosis (**Fig. 6**), groups of mice (n=4-6; 22-24 g) received an intradermal injection of 110 µg west African (Nigeria) *N. nigricollis* venom dissolved in 50 µL PBS. Then, at various time intervals after venom (either immediately [0 min], 15 min, or 60 min), a solution of varespladib (100 µg dissolved in 50 µL PBS) was injected intradermally at the same site of venom injection. In the case of venom only controls, 50 µL of PBS was administered intradermally at the site of venom injection immediately after venom. Similar experiments were then performed using intravenous delivery of varespladib. After the intradermal injection of 110 µg *N. nigricollis* venom, varespladib (100 µg in 50 µL PBS) was administered intravenously in the caudal vein, either immediately (0 min), or 30, 60 or 120 min after venom. Control groups of mice received 110 µg of venom and PBS intravenously instead of varespladib. At 72 h, mice were sacrificed by CO_2_ inhalation, the skin was removed and the areas of the necrotic lesions in the inner side of the skin were measured as previously described.

### *In vivo* models of myotoxicity and treatment with varespladib

To assess inhibition of myotoxicity (**Fig. 6**), west African (Nigeria) *N. nigricollis* venom was incubated with varespladib at 37 °C for 30 minutes. Then, aliquots of 50 µL, each containing 10 µg venom and 10 µg varespladib, were injected intramuscularly into the right gastrocnemius muscle of groups of mice (n=4-5; 18-20 g). Controls included mice receiving 10 µg of venom alone and mice receiving 50 μL PBS alone. Three hours after venom injection, blood samples were collected into heparinised capillary tubes under light inhaled isoflurane anaesthesia by cutting the tip of the tail. After centrifugation, the CK activity of plasma was quantified by using a commercial kit (CK-NAC-UVAA kit; Wiener Laboratories, Rosario, Argentina). CK activity was expressed as units/litre (U/L). Additionally, experiments in which varespladib was administered after venom injection were performed (**Fig. 6**). For this, mice received an intramuscular injection, in the right gastrocnemius, of 10 µg west African (Nigeria) *N. nigricollis* venom, dissolved in 50 µL PBS. At various time intervals after venom injection (either immediately [0 min], or at 15 min), a dose of 100 µg varespladib, dissolved in 50 µL PBS, was administered either intramuscularly at the site of venom injection, or intravenously in the caudal vein. A control group of mice received 10 µg venom intramuscularly and PBS by either route instead of varespladib immediately after venom injection. Another control group received only 50 µL PBS by the intramuscular route.

### Lesion severity scoring using the Venom Induced Dermonecrosis Analysis tool: VIDAL

The severity of the dermonecrotic lesions was assessed using our newly developed AI analyser, VIDAL, the details of which can be found in Laprade, *et al* (*40*). For full details, see **Methods S5**.

### Histopathological analysis of excised tissue samples

Formalin-stored tissue samples were processed and embedded in paraffin, before four micrometer paraffin sections were cut and placed on colour slides or poly-lysine slides to dry. The slides were haematoxylin & eosin stained and cover slipped using DPX. Brightfield images of the H&E-stained lesions were captured with an Echo Revolve microscope, and evidence of necrosis was assessed separately for the epidermis, dermis, hypodermis, panniculus carnosus, and adventitia layers, as described by Hall *et al*. (*41*). The % necrosis of each skin layer within each image was assessed by two independent and blinded pathologists and scored, with mean scores for each layer on each image determined. The ‘dermonecrosis severity score’ was determined for each lesion by taking the mean of the individual layer scores. See **Methods S6** for full methods on the histopathological analysis.

### Statistical analysis

All data are presented as mean average ± standard deviation of at least three independent experimental replicates. Appropriate statistical tests and post-hoc tests were carried out and a difference was considered statistically significant where *P* ≤ 0.05. See **Methods S7** for full details on the statistical analysis.

## Acknowledgments

The authors thank Paul Rowley for maintenance of snakes and provision of venom, Dr. Amy Marriott for assistance with animal welfare observations during in vivo experimentation, Valerie Tilston and colleagues at the University of Liverpool for preparing histopathology slides, and Dr. Matt Lewin for provision of varespladib. The authors acknowledge use of the Biomedical Services Unit provided by Liverpool Shared Research Facilities, Faculty of Health and Life Sciences, University of Liverpool.

## Data and materials availability

All data needed to evaluate the conclusions in the paper are present in the paper, the Supplementary Materials, or via the publicly available FigShare repository: 10.6084/m9.figshare.23580120.

