## Supplementary material for "Dermonecrosis caused by spitting cobra snakebite results from toxin potentiation and is prevented by the repurposed drug varespladib": Fig. S1

**This PDF file includes:**

- Methods S1 to S7
- Figures S1 to S11
- Tables S1 to S3
- Legend for Dataset S1

**Other supporting materials for this manuscript include the following:**

- Dataset S1

**Methods S1: Chemicals, Drugs and Biological Materials**
Thiazolyl blue methyltetrazolium bromide (MTT; M5655), dimethyl sulfoxide (DMSO; 276855), propidium iodide (PI; P4170), and varespladib (SML1100; stock solution of 65.7 mM [25 mg/mL] in DMSO) were purchased from Sigma-Aldrich (Merck). Varespladib used in delayed treatment *in vivo* experiments was provided by Dr. Matthew Lewin (Ophirex Inc., Corte Madera, CA, USA). Dulbecco's modified Eagle's medium (DMEM; 11574516), foetal bovine serum (FBS; 11573397), FluoroBrite DMEM (A1896701), glutaMAX supplement (35050038), penicillin-streptomycin (11528876), phosphate buffered saline (11503387), and TrypLE Express were purchased from Gibco (Thermo Fisher Scientific).

**Methods S2: Venoms**
Venoms were sourced from snake specimens maintained in the Liverpool School of Tropical Medicine's (LSTM) Herpetarium. This facility and its snake husbandry protocols are approved and inspected by the UK Home Office and the LSTM Animal Welfare and Ethical Review Boards. Venom pools were from wild-caught animals of differing geographical origins, namely: *Naja nigricollis* (Tanzania and Nigeria; four individuals in each venom pool), and *N. pallida* (Tanzania; one individual). Crude venoms were lyophilised and stored at 4 °C to ensure long-term stability, then resuspended in PBS (10 mg/mL) and kept at -80 °C until used in the described experiments, with freeze-thaw cycles kept to a minimum to prevent degradation.

**Methods S3: Toxin isolation**
The initial step in toxin isolation was a large-scale size separation on a gel filtration chromatography column; this separates the 3FTx and $PLA_2$ toxins from the higher molecular weight proteins (i.e. snake venom metalloproteinases, L-amino acid oxidase and cysteine rich secretory proteins). For this, 100 mg of *N. nigricollis* (Tanzania) venom was dissolved in 5 mL ice-cold PBS (25 mM sodium phosphate pH 7.2, 0.15 M NaCl) and centrifuged at 10,000 xg for 10 minutes. The supernatant was immediately loaded onto a 480 mL column (2.6 x 95 cm) of Superdex 200HR equilibrated in PBS. The column was operated at a flow rate of 1.0 mL/min during loading and 2.0 mL/min for elution. 10 mL fractions were collected after the void volume and elution was monitored at 280 nm (see **Fig. S1**). SDS-PAGE analysis was carried out on all fractions seen to contain protein on the trace. $PLA_2$ and 3FTx, respectively, were found to be wholly contained in overlapping peaks 1 and 2. Since their separation was incomplete at this stage, these peaks were pooled in preparation for a second step on cation exchange chromatography. This $PLA_2$/3FTx pool was dialysed against 50 mM sodium phosphate, pH 6.0 and then applied to a 20 mL HPSP HiPrep column equilibrated in the same buffer. Elution was carried out using a 15-column volume (CV) gradient of 0 - 0.7 M NaCl in 50 mM sodium phosphate, pH 6.0. The flow rate was 1.5 mL/min. The unbound material was retained and 10 mL fractions collected from the start of the NaCl gradient. Elution was monitored at 280 nm (**Fig. S2**). SDS-PAGE analysis was carried out on the main peaks (**Fig. S3**). Cation exchange peaks 3, 4, and 11 contained a band at 14 kDa, assumed to be $PLA_2$ and based on the relative elution position, 3, 4 are acidic $PLA_2$ and 11 is basic $PLA_2$. Peaks 5-9 each contained equal-sized bands that ran just below the 10 kDa marker, characteristic of 3FTx proteins. The major 3FTx fractions, peaks 6, 8, and 9, were pooled back together and dialysed against PBS to form the whole 3FTx fraction, containing the major 3FTx cytotoxins 1, 3 and 4 [see below], used for the *in vivo* studies, and referred to as CTx in this study.

*Purification of acidic $PLA_2$.* As judged by SDS-PAGE and RP-HPLC (see **Fig. S8**), the $PLA_2$ in peak 4 was sufficiently pure for cytotoxicity studies. It was found to have a mass of 13,287 Da. MS/MS of tryptic peptides (see methods below) matched this protein to acidic phospholipase A2 CM-I P00602 from *N. mossambica* (see **Table S1**). The $PLA_2$ in peak 3 required a third chromatography step for full purity. This fraction was loaded directly onto a 4.7 mL column of Phenyl Sepharose LS FF equilibrated in 25 mM sodium phosphate pH 7.2. Elution was carried out with a 0-100% (4 CV) gradient of 25 mM sodium phosphate pH 7.2 to 25% ethanol in 25 mM sodium phosphate pH 7.2 with long hold step (5 CV) in the latter buffer. The column was operated at 1.0 mL/min and elution was monitored at 214 nm (see **Fig. S4**). The mass of the purified protein, eluting between 30 and 40 minutes, was found to be 13,172 Da and it was also matched to acidic phospholipase A2 CM-I P00602 ('acidic $PLA_2$') from *N. mossambica* following trypsin digestion and MS/MS (see **Table S1**). The two acidic $PLA_2$ forms were combined and used as a single acidic $PLA_2$ pool throughout the study.

*Purification of basic $PLA_2$.* Peak 11 from the cation exchange chromatography step was dialysed against 5 mM sodium phosphate pH 6.8 and then loaded onto a 1 mL column of ceramic hydroxyapatite (CHT I, BioRad) equilibrated in the same buffer. Using a flow rate of 0.5 mL/min, elution was carried out with gradient of 5 mM sodium phosphate pH 6.8 to 500 mM sodium phosphate pH 6.8, 0.15 M NaCl over 20 CV. Elution was monitored at 214 nm (see **Fig. S5**). The protein in the main peak (eluting between 35 and 40 mins) was pure as determined by SDS-PAGE and RP-HPLC (see **Fig. S8**), and was found to have a mass of 13,249 Da. Trypsin digestion with MS/MS analysis resulted in a match to Phospholipase A2 'basic' P00605 ('basic $PLA_2$') from *N. nigricollis* (see **Table S1**).

The basic $PLA_2$ and the two acidic $PLA_2$ were pooled back together and dialysed against PBS to form the combined $PLA_2$ fraction used in the *in vivo* studies.

*Purification of individual 3FTx cytotoxins for cell cytotoxicity studies.* The material in peak 6 from cation exchange chromatography was fairly pure as determined by RP-HPLC, but it was highly purified using hydrophobic interaction chromatography (HIC). For this, the fraction was made up to 1.5 M in NaCl and loaded onto a 10 mL Phenyl Superose column. Proteins were then eluted in a 5 CV gradient of 1.5 M NaCl in 25 mM sodium phosphate pH 7.2 to 30% (v/v) ethylene glycol in 25 mM sodium phosphate pH 7.2. The flow rate was 1 mL/min and elution was monitored at 214 nm (see **Fig. S6**). The protein in the main peak (eluting at 50 mins) was found to be pure by SDS-PAGE and RP-HPLC and its mass was determined to be 6,817 Da. MS/MS of tryptic peptides confirmed this protein to be Cytotoxin 1 P01468, referred to as CTx1 throughout this study.

RP-HPLC analysis showed that peaks 8 and 9 together contained three main 3FTx forms. These were separated using HIC as for peak 6. The conditions were identical, except that the starting buffer was 1.2 M NaCl rather than 1.5 M. As can be seen in **Fig. S7**, the three proteins are fully separated by these means. RP-HPLC analysis showed each of these to be pure (**Fig. S9**) and to possess very characteristic peak shapes. All three were subjected to digestion with trypsin and identified using MS/MS. Peak 12 had a mass of 6,707 Da and was identified to be Cytotoxin 4 P01452 (CTx4 in this study). Peak 13 had a mass of 6,817 Da and was identified using MS/MS to be Cytotoxin 1 P01468. This is likely to be a version of the protein identified in peak 6 with a Asn/Asp modification and is named hereafter as CTx1v. Peak 14 had a mass of 6,884 Da and was determined to be Naniproin/Cytotoxin 3 P0DSN1 (CTx3).

*Purity analysis methods.* SDS-PAGE was carried out using 4-20% acrylamide gels (BioRad) and run using a Tris-glycine buffer system, followed by staining with Coomassie Blue R250. RP-HPLC was performed on a Vanquish HPLC system (Thermo) using a Biobasic C4 column (2.1 x 150 mm). The flow rate was 0.2 mL/min and proteins were separated in a gradient of acetonitrile (see **Fig. S8** and **S9** for gradient formats) in 0.1% trifluoroacetic acid, with monitoring at 214 nm. Between 2-5 µg protein was typically loaded per analysis.

*Trypsin digestion and MS/MS analysis.* In preparation for analysis, the relevant proteins were desalted on a RP-HPLC column, as above. These were then dried in a centrifugal evaporator, 20 µL $H_2O$ was added and then re-dried. The proteins were then resuspended in 8 M urea/0.1 M Tris-Cl (pH 8.5), reduced with 5 mM TCEP (tris (2-carboxyethyl) phosphine) for 20 min and alkylated with 50 mM 2-chloroacetamide for 15 min in the dark at room temperature. Samples were diluted 4-fold with 100 mM Tris-Cl (pH 8.5) and digested with trypsin at an enzyme/substrate ratio of 1:20 overnight at 37°C. The reaction was terminated by addition of formic acid (FA), and digested peptides were loaded on to Evotips and analysed directly using an Evosep One liquid chromatography system (Evosep Biosystems, Denmark) coupled with timsTOF SCP-mass spectrometer (Bruker, Germany). Peptides were separated on a 75 µm i.d. × 15 cm separation column packed with 1.9 µm C18 beads (Evosep Biosystems, Denmark) and over a predetermined 44-minute gradient. Buffer A was 0.1% FA in water and buffer B was 0.1% FA in acetonitrile. Instrument control and data acquisition were performed using Compass Hystar (version 6.0) with the timsTOF SCP operating in data-dependent acquisition mode. Fragmentation spectra were searched against an in-house venom gland-derived protein sequence database using Mascot (*21, 77*). Reverse decoys and contaminants were included in the search database. Cysteine carbamidomethylation was selected as a fixed modification, and oxidation of methionine was selected as a variable modification. The precursor-ion mass tolerance and fragment-ion mass tolerance were set at 10 ppm and 0.04 Da, respectively, and up to two missed tryptic cleavages were allowed. Mascot files were parsed into Scaffold (version 5.0.1, Proteome Software, Inc.) for validation at a protein-level false discovery rate (FDR) of <1%.

**Methods S4: Animal ethics and maintenance**

Animal experiment protocols were performed using approvals from the Animal Welfare and Ethical Review Boards of the Liverpool School of Tropical Medicine and the University of Liverpool, with licensed approval (PPL# P58464F90) from the UK Home Office, and in accordance with the UK Animal (Scientific Procedures) Act 1986, and the Institutional Committee for the Care and Use of Laboratory Animals (CICUA) of the University of Costa Rica (approval number CICUA 82-2). Male SWISS (CD1) mice (18-27 g; Charles River, Janvier, Animal Facility of Instituto Clodomiro Picado, Costa Rica) were housed in groups of five and acclimated for one week prior to experimentation. Mice used in experiments conducted in Liverpool (**Fig. 3-5**) were given *ad libitum* access to CRM irradiated food (Special Diet Services, UK) and reverse osmosis water in an automatic water system and kept under room conditions of approximately 22 °C, 40–50% humidity, with 12/12-hour light cycles. Mice used in experiments conducted in Costa Rica (**Fig. 1, 6**) were maintained under conditions of 22-24 °C and 60-65% humidity, with 12/12-hour light cycles, and given *ad libitum* access to food and water.

**Methods S5: Lesion severity scoring using the Venom Induced Dermonecrosis Analysis tool: VIDAL**

The severity of the dermonecrotic lesions was assessed using our newly developed AI analyser, VIDAL, the details of which can be found in Laprade, *et al* (1). Briefly, images of the dissected lesions on A4 standardisation cut out masks (2) were uploaded to the AI's website (https://github.com/laprade117/VIDAL-Experiments). The program measured and scored the dark and light regions of the lesions from which it calculated a total dermonecrosis score, given the appropriately named Dermonecrotic Units (DnU), where the higher the number the more severe the lesion.

**Methods S6: Histopathological analysis of excised tissue samples**

Formalin-stored tissue samples were processed using a Tissue-Tek VIP (vacuum infiltration processor) overnight, before being embedded in paraffin (Ultraplast premium embedding medium, Solmedia, WAX060). Four micrometer paraffin sections were cut on a Leica RM2125 RT microtome, floated on a water bath, and placed on colour slides (Solmedia, MSS54511YW) or poly-lysine slides (Solmedia MSS61012S) to dry. For haematoxylin & eosin (H&E) staining, slides were dewaxed in xylene and rehydrated through descending grades of ethanol (100%, 96%, 85%, 70%) to distilled water before being stained in haematoxylin (5 mins), “blued” in tap water (5 mins) and stained in eosin (2 mins). Slides were then dehydrated through 96% and 100% ethanol to xylene and cover slipped using DPX (CellPath SEA-1304-00A). Haematoxylin (Atom Scientific, RRBD61-X) and eosin (TCS, HS250) solutions were created in house. Brightfield images of the H&E-stained lesions were captured with an Echo Revolve microscope (Settings: 10x magnification; LED: 100%; Brightness: 30; Contrast: 50; Colour balance: 50), with at least five images taken per slide. As described by Hall *et al*. (3), histological evidence of necrosis was assessed separately for the epidermis, dermis, hypodermis, panniculus carnosus, and adventitia layers. Features of necrosis included loss of nuclei, nuclear fragmentation (karyorrhexis), nuclear shrinkage and hyperchromasia (pyknosis), loss of cytoplasmic detail with hypereosinophilia, loss of cell borders and, in the case of severe necrosis, disarray with complete loss of architecture and hyalinization. In the epidermis, ulceration with superficial debris was interpreted as evidence of necrosis. In the dermis, loss of skin adnexal structures (e.g., hair follicles and sebaceous glands) and extracellular matrix disarray were also interpreted as evidence of necrosis. The %-necrosis of each skin layer (epidermis, dermis, hypodermis, panniculus carnosus, and adventitia) within each image was assessed by two independent and blinded pathologists and scored between 0 and 4, with 0 meaning no observable necrosis in the layer within that image, 1 meaning up to 25% of the layer in that image exhibiting signs of necrosis, 2 meaning 25-50% necrosis, 3 meaning 50-75%, and 4 meaning more than 75% exhibiting indicators of necrosis. The mean scores of the pathologists for each layer from each image were determined, and the highest scores-per-mouse used for our data analysis as these represented the maximum necrotic severity within each lesion. The ‘dermonecrosis severity score’ was determined for each lesion by taking the mean of the individual layer scores.

**Methods S7: Statistical analysis**

All data are presented as mean average ± standard deviation of at least three independent experimental replicates. For cell experiments, ‘n’ is defined as an independent experiment completed at a separate time from other ‘n’s within that group of experiments; all drug and/or venom treatments within an ‘n’ were completed in triplicate wells and the mean taken as the final value for that one trial. For *in vivo* experiments, ‘n’ is defined as the number of mice in that specific treatment group. Two-tailed t-tests were performed for dual comparisons, one-way analysis of variances (ANOVAs) performed for multiple comparisons with one independent variable followed by Dunnett’s or Tukey’s multiple comparisons tests when the trial data were compared to a single control group or to all other groups, respectively, as recommended by GraphPad Prism, and two-way ANOVAs performed for multiple comparisons with two independent variables followed by Dunnett’s multiple comparisons tests. A difference was considered statistically significant where $P \leq 0.05$.

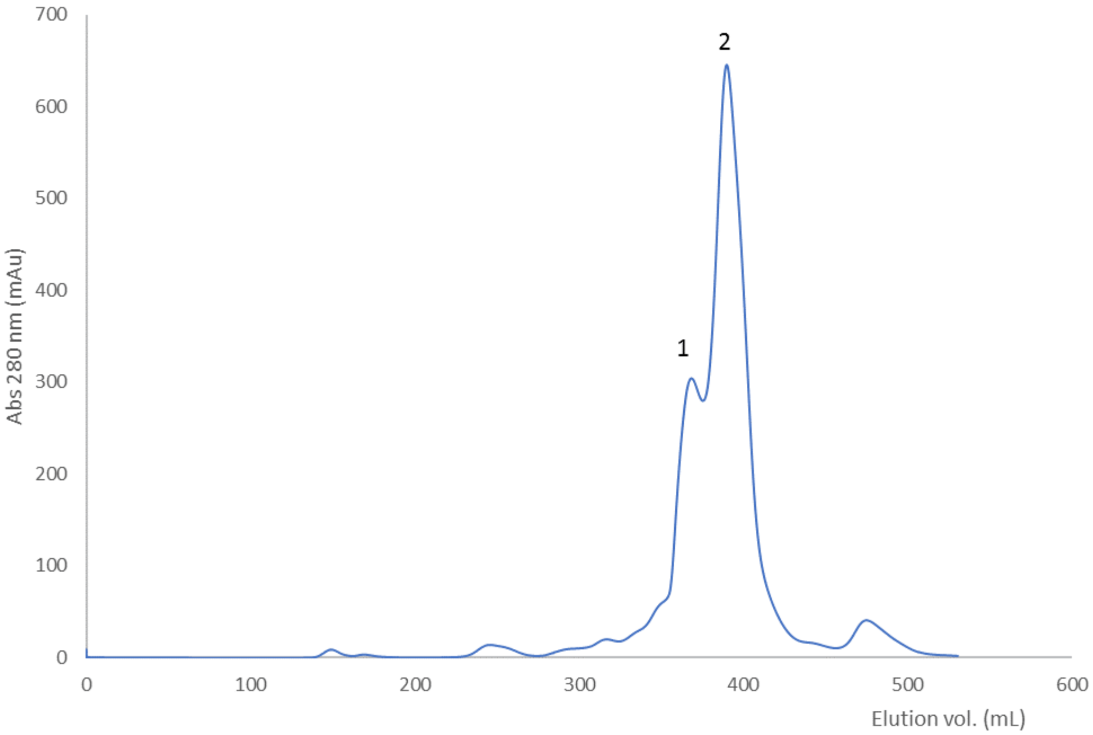

**Fig. S1. Gel filtration chromatography of whole venom.**

100 mg of *N. nigricollis* (Tanzania) venom was dissolved in 5 mL PBS and centrifuged prior to separation. This was loaded onto a 480 mL column of Superdex 200HR equilibrated in PBS. The column was operated at a flow rate of 2.0 mL/min and elution was monitored at 280 nm. The overlapping peaks for $PLA_2$ and 3FTX are numbered 1 and 2 respectively. These were pooled and then subjected to cation exchange chromatography (see Fig. S2).

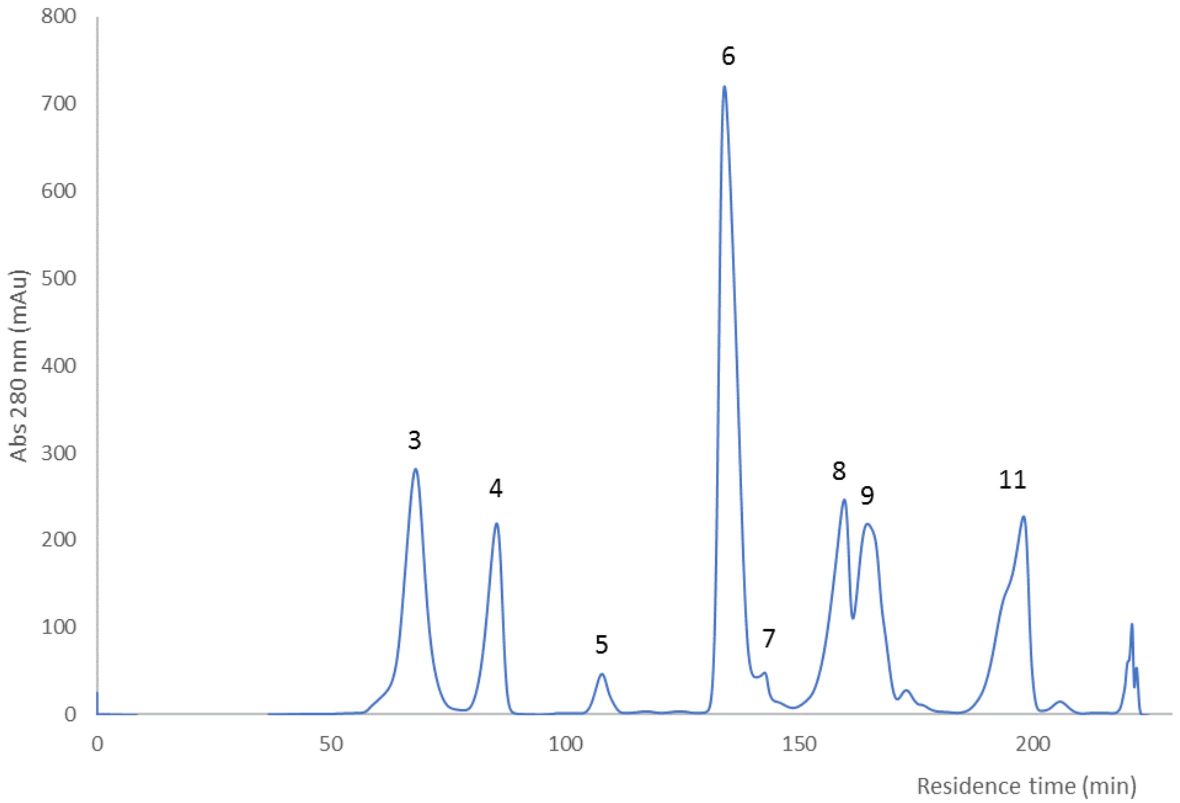

**Fig. S2. Cation exchange chromatography of the $PLA_2$ and 3FTX fraction from GFC.**
Peaks 1 and 2 from GFC were pooled and dialysed against 50 mM sodium phosphate, pH 6.0 and then applied to a 20 mL HPSP column equilibrated in the same buffer. Elution was carried using a 15 CV gradient of 0 - 0.7 M NaCl in 50 mM sodium phosphate, pH 6.0. Flow rate was 1.5 mL/min and elution was monitored at 280 nm. Peak 4 contained pure acidic $PLA_2$ (named acidic $PLA_2$ 2). The major peaks, labelled 3, 6, 8, 9, and 11 were subject to further chromatographic steps.

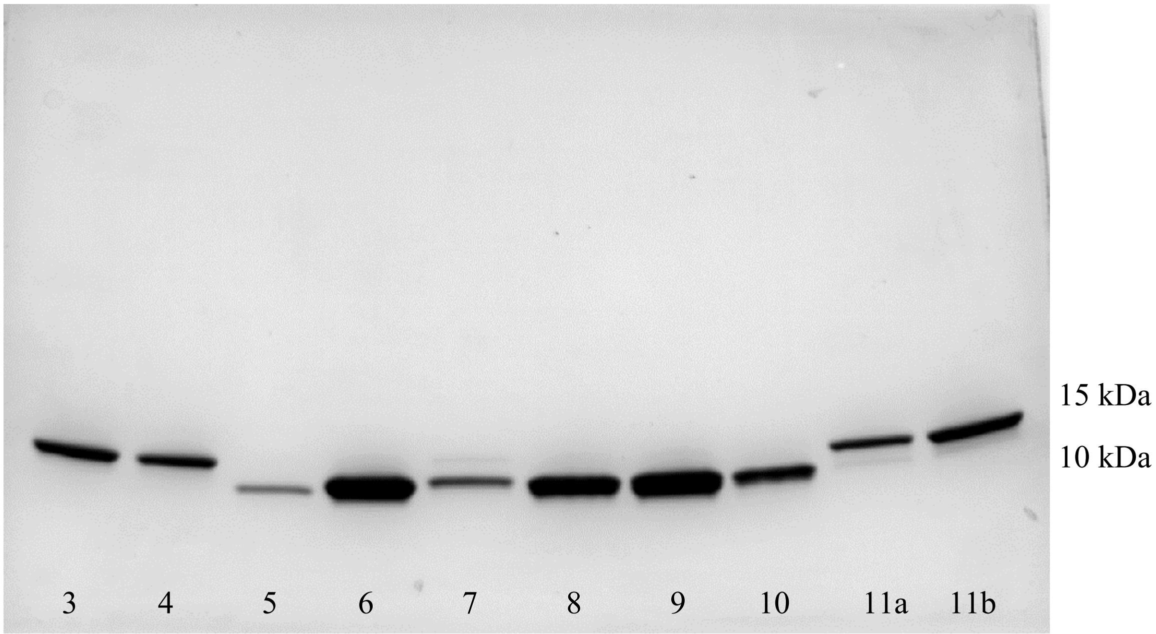

**Fig. S3. SDS-PAGE analysis of peaks from cation exchange chromatography.**

Seven µL samples were loaded onto 4-20% acrylamide gels (BioRad) and run using the Tris-glycine buffer system under reducing conditions. The gels were subsequently stained with Coomassie Blue R250. The position of relevant molecular weight markers is shown on the right (kDa). Chromatography peak numbers are indicated at the base of the figure. Peak 11 was run as two fractions, a and b, reflecting the wide asymmetric peak shape. Peak 10 contained overlapping material from peak 9, hence the band intensity is greater than would be expected for the height of peak 10.

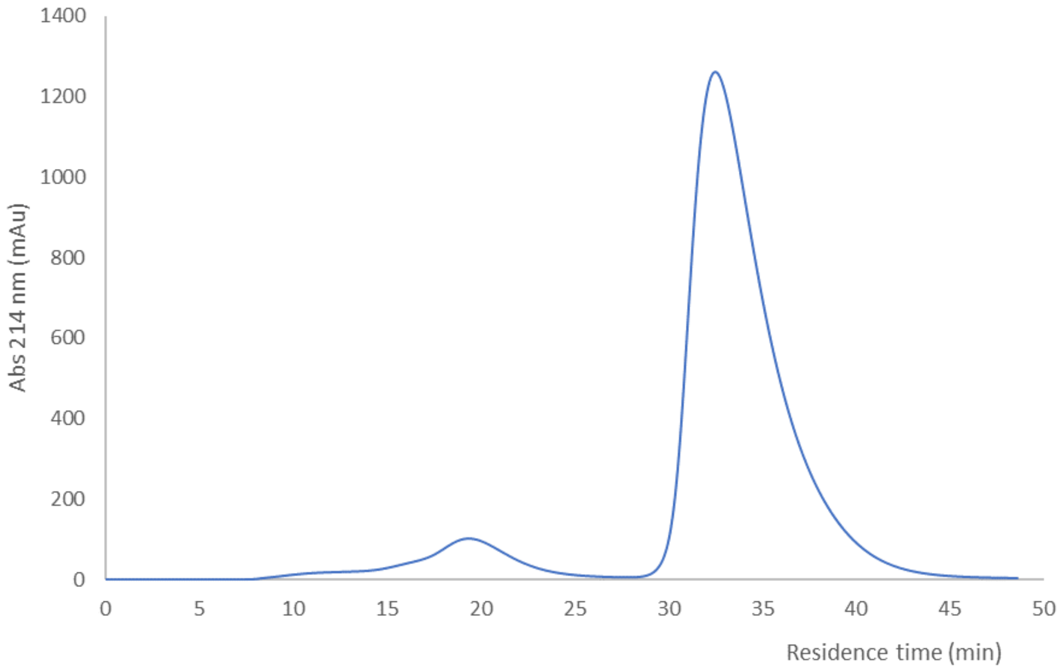

**Fig. S4. Hydrophobic interaction chromatography of acidic $PLA_2$ (peak 3).**
Peak 3 from the cation exchange chromatography step was loaded directly onto a 4.7 mL column of Phenyl Sepharose LS FF equilibrated in 25 mM sodium phosphate pH 7.2. Elution was carried out with a 0-100% (4CV) gradient of 25 mM sodium phosphate pH 7.2 to 25% ethanol in 25 mM sodium phosphate pH 7.2. The column was operated at 1.0 mL/min and elution was monitored at 214 nm.

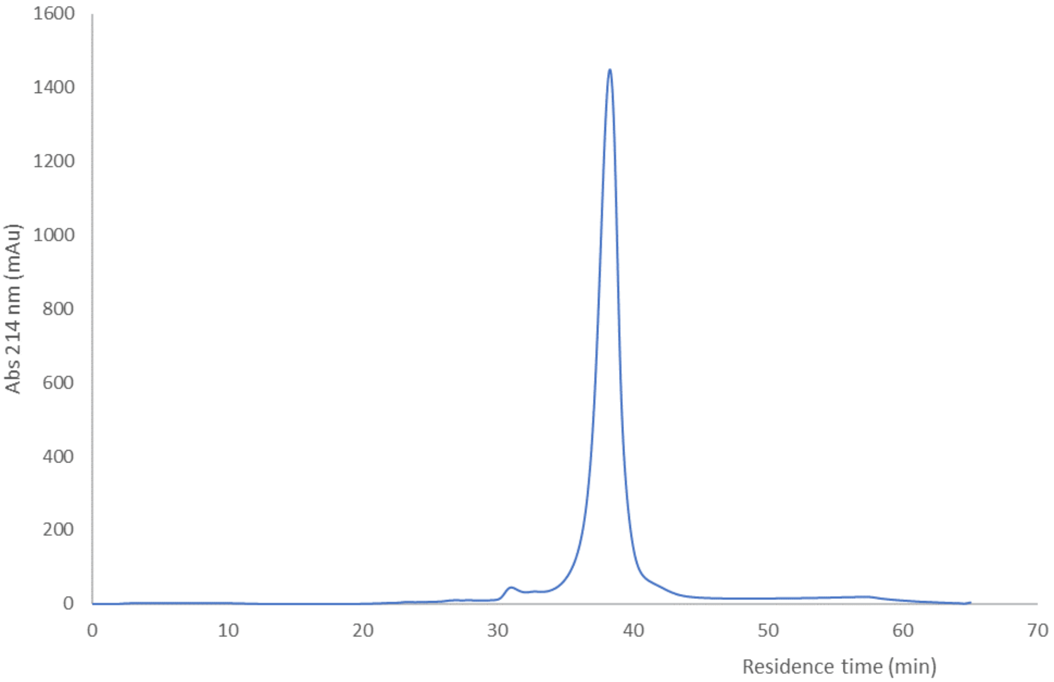

**Fig. S5. Hydroxyapatite chromatography of basic $PLA_2$ (peak 11).**

Peak 11 from the cation exchange chromatography step was dialysed against 5 mM sodium phosphate pH 6.8 and then loaded onto a 1 mL column of ceramic hydroxyapatite (CHT I) equilibrated in the same buffer. Using a flow rate of 0.5 mL/min, elution was carried out with gradient of 5 mM sodium phosphate pH 6.8 to 500 mM sodium phosphate pH 6.8, 0.15 M NaCl over 20 CV. Elution was monitored at 214 nm.

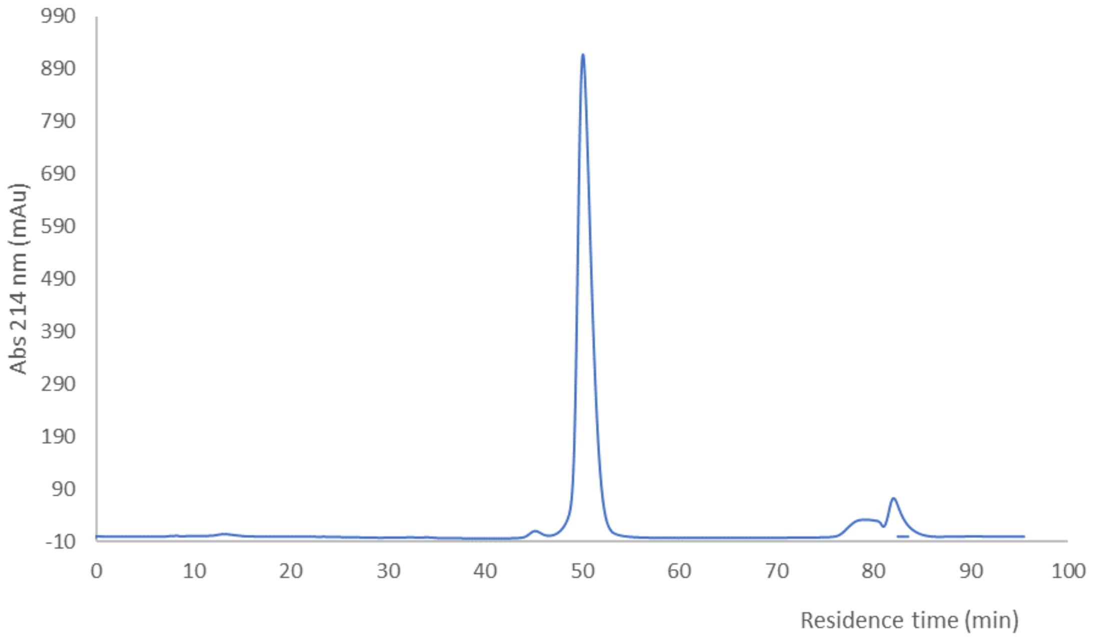

**Fig. S6. Hydrophobic interaction chromatography of peak 6 3FTx cytotoxin 1.**

The material in peak 6 from cation exchange chromatography was made to 1.5 M in NaCl and loaded onto a 10 mL Phenyl Superose column. Proteins were then eluted in a 5 CV gradient of 1.5 M NaCl in 25 mM sodium phosphate pH 7.2 to 30% (v/v) ethylene glycol in 25 mM sodium phosphate pH 7.2. The flow rate was 1 mL/min and elution was monitored at 214 nm.

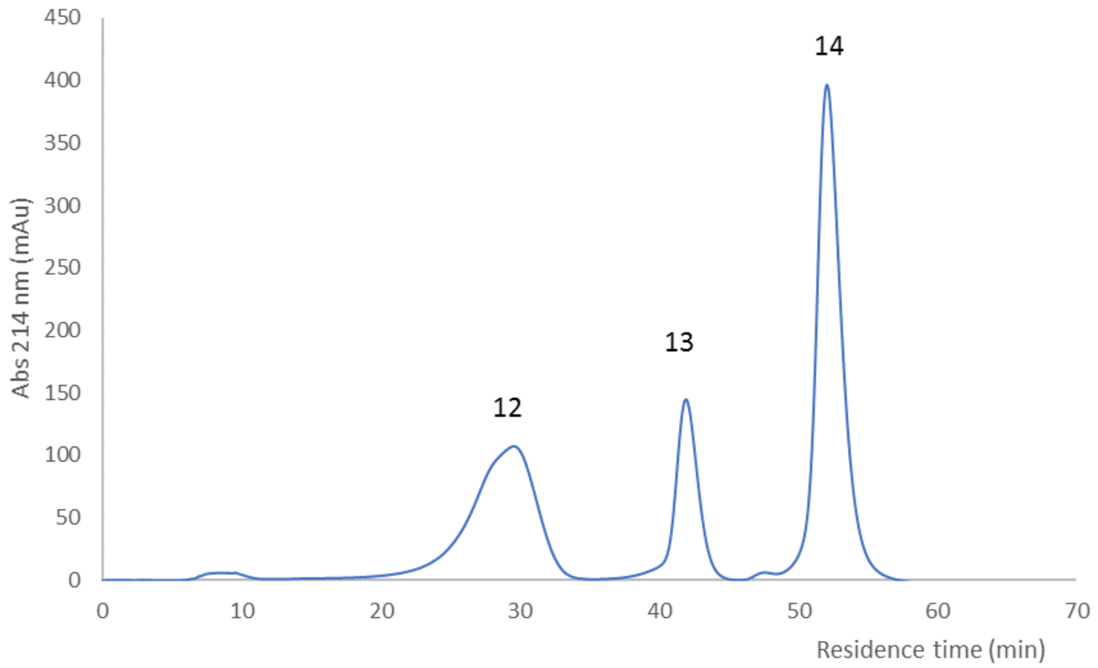

**Fig. S7. Hydrophobic interaction chromatography of 3FTx cytotoxins 1v, 3 and 4.**
The material in peak 8 and 9 from cation exchange chromatography was made to 1.2 M in NaCl and loaded onto a 10 mL Phenyl Superose column. Proteins were then eluted in a 5 CV gradient of 1.2 M NaCl in 25 mM sodium phosphate pH 7.2 to 30% (v/v) ethylene glycol in 25 mM sodium phosphate pH 7.2. The flow rate was 1 mL/min and elution was monitored at 214 nm. Peak 12, 13 and 14 respectively contained cytotoxin 4, cytotoxin 1v and cytotoxin 3.

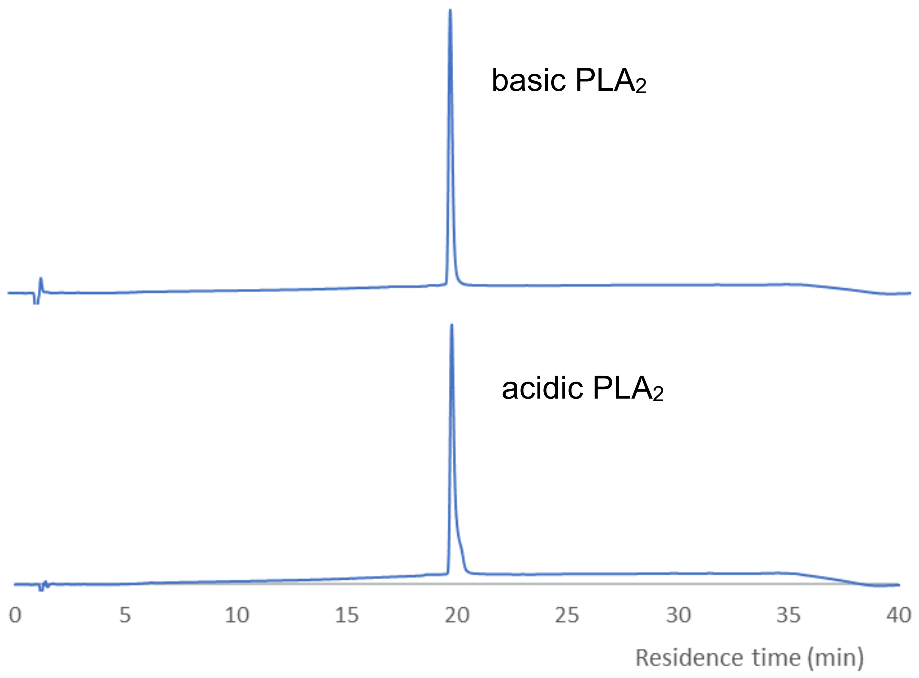

**Fig. S8. Analytical RP-HPLC of purified $PLA_2$s.**

RP-HPLC was performed on a Biobasic C4 column (2.1 x 150 mm). The flow rate was 0.2 mL/min and proteins were separated in a gradient of 0-65% acetonitrile in 0.1% trifluoroacetic acid, with monitoring at 214 nm (Y-axis, mAu units not shown). The acidic $PLA_2$ shown is acidic $PLA_2$ 1; acidic $PLA_2$ 2 had an identical profile on RP-HPLC and the two were combined to give the acidic $PLA_2$ pool used throughout this study.

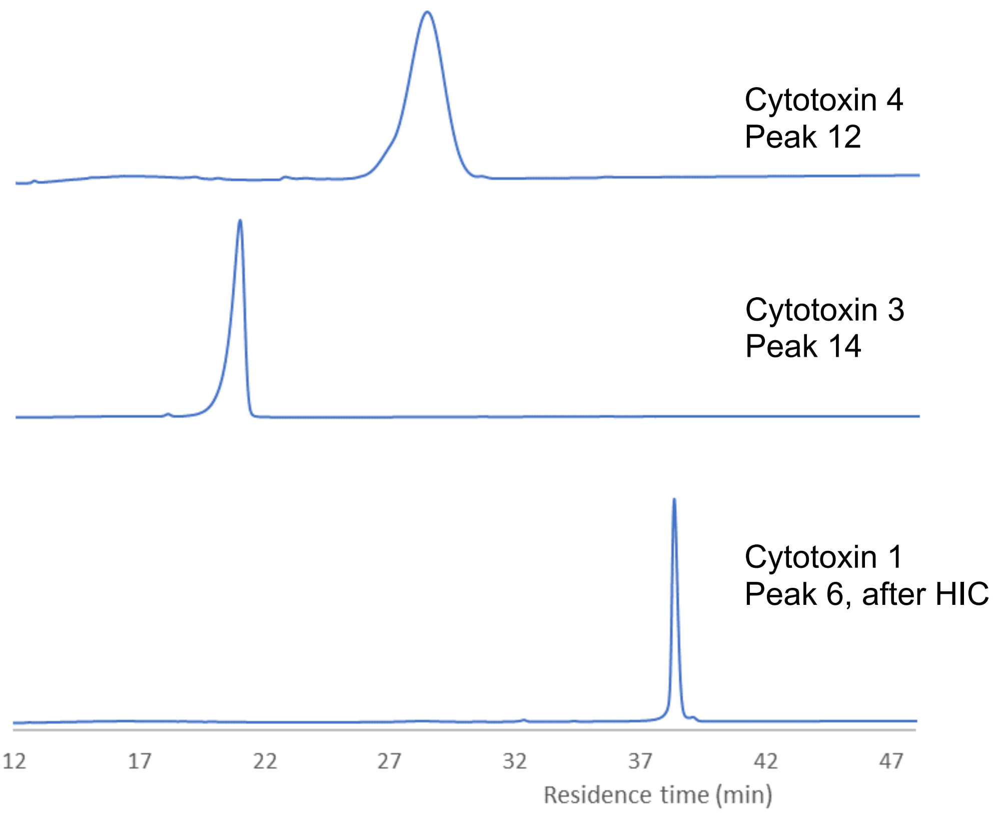

**Fig. S9. Analytical RP-HPLC of purified 3FTx cytotoxins 1, 3 and 4.**
RP-HPLC was performed on a Biobasic C4 column (2.1 x 150 mm). The flow rate was 0.2 mL/min and proteins were separated in a gradient of acetonitrile (0-25%/5 min, 25-50%/30 min, 50-90%/5 min) in 0.1% trifluoroacetic acid, with monitoring at 214 nm (Y-axis, mAu units not shown).

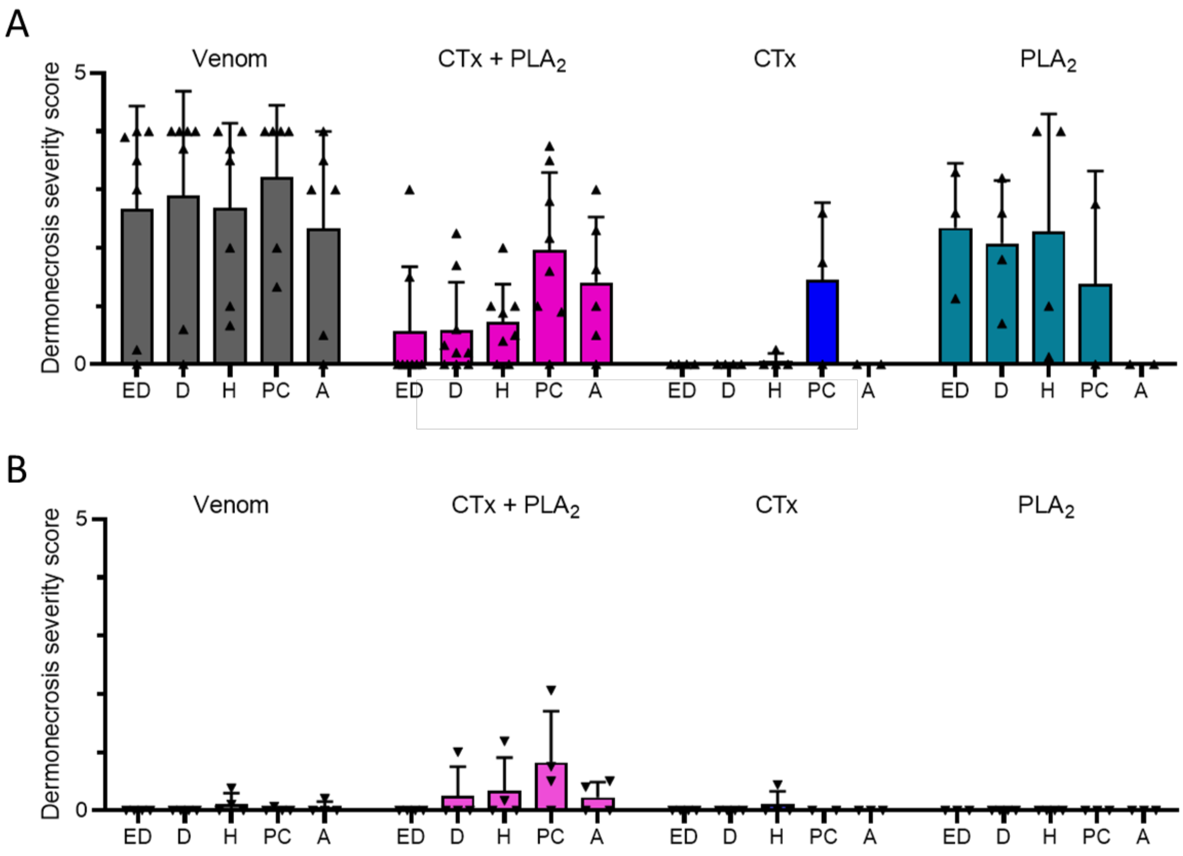

**Fig. S10. Histopathological analysis of dermonecrotic skin lesions shows that microscopic pathology caused by *N. nigricollis* venom is prevented by the $PLA_2$ inhibitor varespladib.**
**A**) Histopathology analysis resulting in dermonecrosis severity scores for multiple skin layers are shown following intradermal dosing of mice with crude East African (Tanzania) *N. nigricollis* venom, the CTX + $PLA_2$ combination, and CTXs and $PLA_2$s separately. Key: ED, epidermis; D, dermis; H, hypodermis; PC, panniculus carnosus; A, adventitia. **B**) Co-injection of mice with the $PLA_2$ inhibitor varespladib significantly reduces damage scores across multiple skin layers. Data shown represents the mean damage score for each skin layer and error bars represent the corresponding standard deviations.

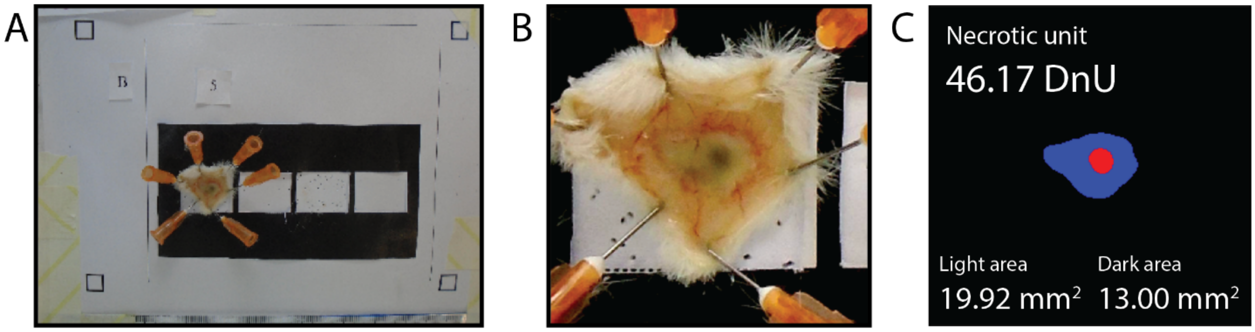

**Fig. S11. Overview of the VIDAL workflow.** First, **A**) the raw image is imported and automatically standardised. Thereafter, **B**) the lesion is identified and cropped out for further analysis. Finally, **C** the processed image is segmented and light and dark lesions automatically identified. This information is used to calclulate dermonecrotic units (DnUs) for each identified lesion, with a 2:1 weighting of dark to light lesion areas.

**Table S1. Summary of MS analysis and identification of the purified toxins.**

| Chromatography peak no. | | Intact mass | MS/MS ID | | | | | |
|---|---|---|---|---|---|---|---|---|
| CatX | HIC | (Mono, Da) | Protein ID | Entry Name | Length | Percent Coverage | Organism | Protein Description |
| 3 | | 13,172 | P00602 | PA2A1_NAJMO | 118 | 85.6 | Naja mossambica OX=8644 | Acidic phospholipase A2 CM-I |
| 4 | | 13,287 | P00602 | PA2A1_NAJMO | 118 | 85.6 | Naja mossambica OX=8644 | Acidic phospholipase A2 CM-I |
| 11 | | 13,249 | P00605 | PA2B4_NAJNG | 118 | 93.2 | Naja nigricollis OX=8654 | Phospholipase A2 'basic' |
| 6 | | 6,818 | P01468 | 3SA1_NAJPA | 60 | 100 | Naja pallida OX=8658 | Cytotoxin 1 |
| 8+9 | 12 | 6,707 | P01452 | 3SA4_NAJMO | 60 | 63.3 | Naja mossambica OX=8644 | Cytotoxin 4 |
| 8+9 | 13 | 6,817 | P01468 | 3SA1_NAJPA | 60 | 100 | Naja pallida OX=8658 | Cytotoxin 1 |
| 8+9 | 14 | 6,884 | P0DSN1 | 3SAN_NAJNG | 60 | 96.7 | Naja nigricollis OX=8654 | Naniproin [cytotoxin 3] |

**Table S2. Dermonecrotic lesions from mice dosed with *N. nigricollis* venom and toxins, with and without varespladib.**

Mice were intradermally injected with East African (Tanzania) *N. nigricollis* venom (63 µg), or proportional amounts of purified CTx + $PLA_2$s (37.8 + 16.4 µg), purified CTx-only (37.8 µg), and purified $PLA_2$s-only (16.4 µg) either alone, or pre-incubated with the $PLA_2$-inhibitor varespladib (19 µg). Mice were culled at 72 h post-injection, lesions excised, and photographs taken. White scale bars represent 5 mm.

| | |
|---|---|
| **Venom** | 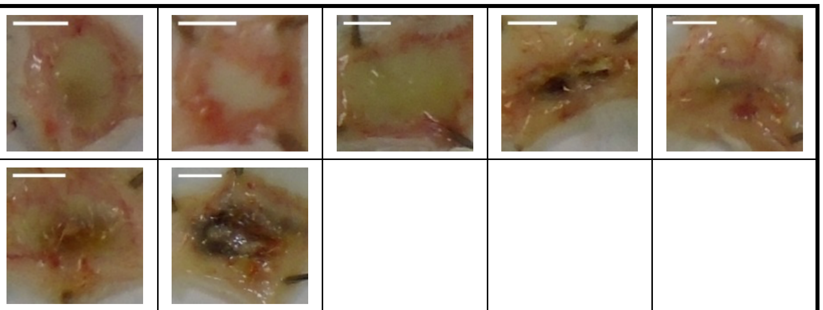 |
| **Venom + varespladib** | 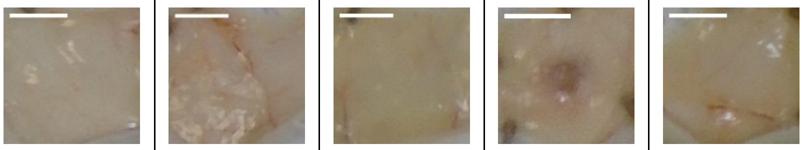 |
| **CTXs + $PLA_2$s** | 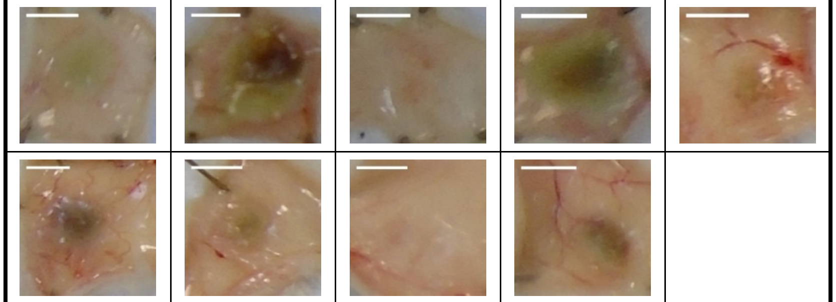 |
| **CTXs + $PLA_2$s + varespladib** | 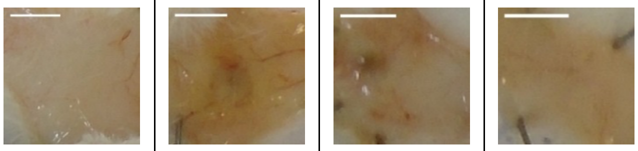 |
| **CTXs-only** | 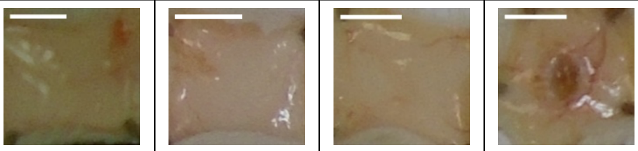 |
| **CTXs-only + varespladib** | 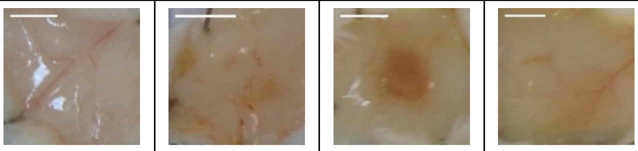 |
| **$PLA_2$s-only** | 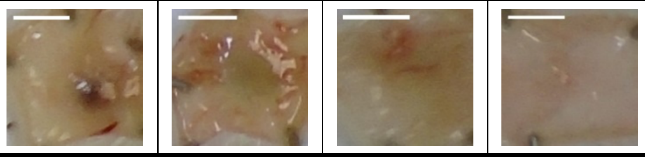 |

| | | | | |
|---|---|---|---|---|
| **$PLA_2$s + varespladib** | 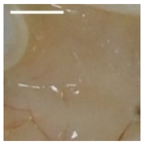 | 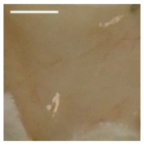 | 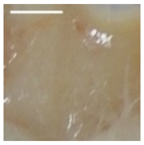 | 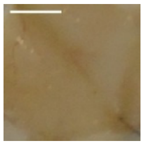 |

**Table S3. Dermonecrotic lesions from mice dosed with *N. pallida* venom, with and without varespladib.**

Mice were intradermally injected with *N. pallida* venom (25 µg) either alone, or pre-incubated with the $PLA_2$-inhibitor varespladib (19 µg). Mice were culled at 72 h post-injection, lesions excised, and photographs taken. White scale bars represent 5 mm.

| | | | | | |
|---|---|---|---|---|---|
| **Venom** | 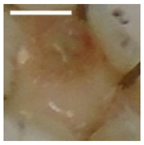 | 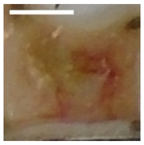 | 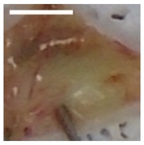 | 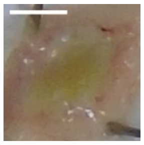 | |
| **Venom + varespladib** | 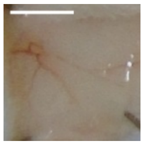 | 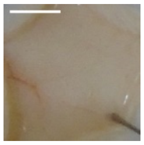 | 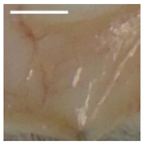 | 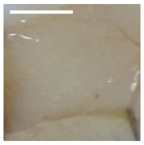 | 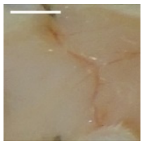 |

**Data S1. All data used for figures presented in the manuscript and SI file.** Data for each figure is contained within an individually labeled sheet within this data file, with individual panels labelled where appropriate.
